# Discovery of antimicrobial compounds from *Lendenfeldia*, *Ircinia* and *Dysidea* sponges using bioassay guided fractionation of marine extracts

**DOI:** 10.1101/489849

**Authors:** Mojdeh Dinarvand, Nicholas Proschogo, Malcolm P. Spain, Gayathri Nagalingam, James A. Triccas, Peter J. Rutledge

## Abstract

Multidrug resistant bacteria have emerged as a threat to public health all over the world. At the same time, the discovery of new bioactive small molecules with antimicrobial activity and suitable pharmacological properties has waned. Herein we report the screening of marine extracts to identify novel compounds with antimicrobial activity. Bioassay guided fractionation has enabled the discovery and identification of a family of simple amines with promising activity against methicillin resistant *Staphylococcus aureus* (MRSA). To confirm the natural product structures proposed, these compounds and analogues have been prepared synthetically. Several of the synthetic analogues showed promising bioactivity against the medically important pathogens MRSA (MICs to 12.5 µM), *Mycobacterium tuberculosis* (MICs to 0.02 µM), uropathogenic *Escherichia coli* (MIC 6.2 µM) and *Pseudomonas aeruginosa* (MIC 3.1 µM). Cross-referencing antimicrobial activity and toxicity show that these synthetic compounds display a favourable therapeutic index for their target pathogens.

## 1. INTRODUCTION

Marine ecosystems have long been a rich source of bioactive natural products, in the search for interesting molecules and novel therapeutic agents.^1–5^ Many interesting and structurally diverse secondary metabolites have been isolated from marine sources and characterised over the last 70 years.^6–9^ Yet the first ‘drugs from the sea’ were only approved in the early 2000s: the cone snail peptide ziconotide (ω-conotoxin MVIIA) in 2004 to alleviate chronic pain,^10^ and sea squirt metabolite trabectedin in 2007 for treatment of soft-tissue sarcoma.^11^ Interest in marine natural products has continued to grow since,^6, 8–9^ spurred in part by the spread of antimicrobial resistant pathogens and the need for new drugs to combat them.^3^

Human pathogens are associated with a variety of moderate to severe infections and the recent rise of multi-drug resistant pathogens makes treatment more difficult. The last two decades have seen the emergence of methicillin resistant *Staphylococcus aureus* (MRSA) strains resistant even to ‘drugs of last resort’ such as vancomycin,^12^ and *Mycobacterium tuberculosis* resistant to all front-line drugs,^13–15^ which highlights the urgent need to find new effective antibiotics. Natural products continue to offer a productive source of structural diversity and bioactivity, and are an important source for new drugs.^3–4, 6–7^

In the search for new antimicrobial agents, we screened a set of marine extracts ^16^ to determine activity against antibiotic resistant microorganisms using a high-throughput screening (HTS) assay. Fractionation and purification of active components by high-performance liquid chromatography (HPLC) and structural elucidation using high resolution and tandem mass spectrometry (MS) led us to a series of potential structures for new, bioactive amine natural products (Figure 1).

To validate the proposed structures, and to explore the potential of this compound class more broadly, analogues based on the general structures **1–5** were synthesised and evaluated as antimicrobial agents against a panel of medicinally important microorganisms: MRSA, *M. tuberculosis*, uropathogenic *Escherichia coli*, and *Pseudomonas aeruginosa*.

**Figure 1.**
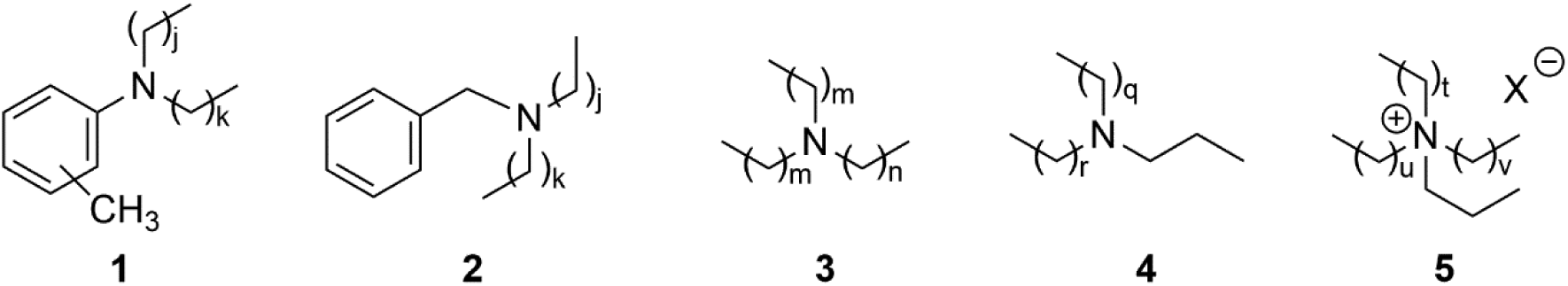
Proposed structures of bioactive amine natural products identified as lead compounds in this study; (j + k) = 14; m = 5, n = 9; (q + r) = 20; (t + u + v) = 19; X = unidentified counterion.

## 2. RESULTS AND DISCUSSION

### 2.1. Identification of Active Marine Extracts

To identify marine samples with activity against MRSA, 1434 compounds from the AIMS Bioresources Library^16^ (provided by the Queensland Compound Library,^17^ now called Compounds Australia^18^) were screened in a resazurin cell viability assay. Of the samples tested, 29 inhibited the growth of MRSA by greater than 50% compared to non-treated controls (Figure S1, Supporting Information). Minimum inhibitory concentrations (MICs) were determined for the 23 most promising samples, representing extracts and fractions from the phyla Porifera (90%), Echinodermata (5%) and Chordata (5% (Table 1 and Table S1). The five most active samples showed MICs at 31.3 µg mL^−1^ (all Porifera samples), while another four samples returned MICs of 62.5 µg mL^−1^ (also all Porifera).

Cytotoxicity screens against HepG2, HEK 293, A549 and THP-1 cell lines were performed to define the cytotoxicity profile of the most active samples. Pleasingly, all the samples most active against MRSA were also nontoxic to the cell lines tested (Tables 1, S1, S2 and S3).

**Table 1.**
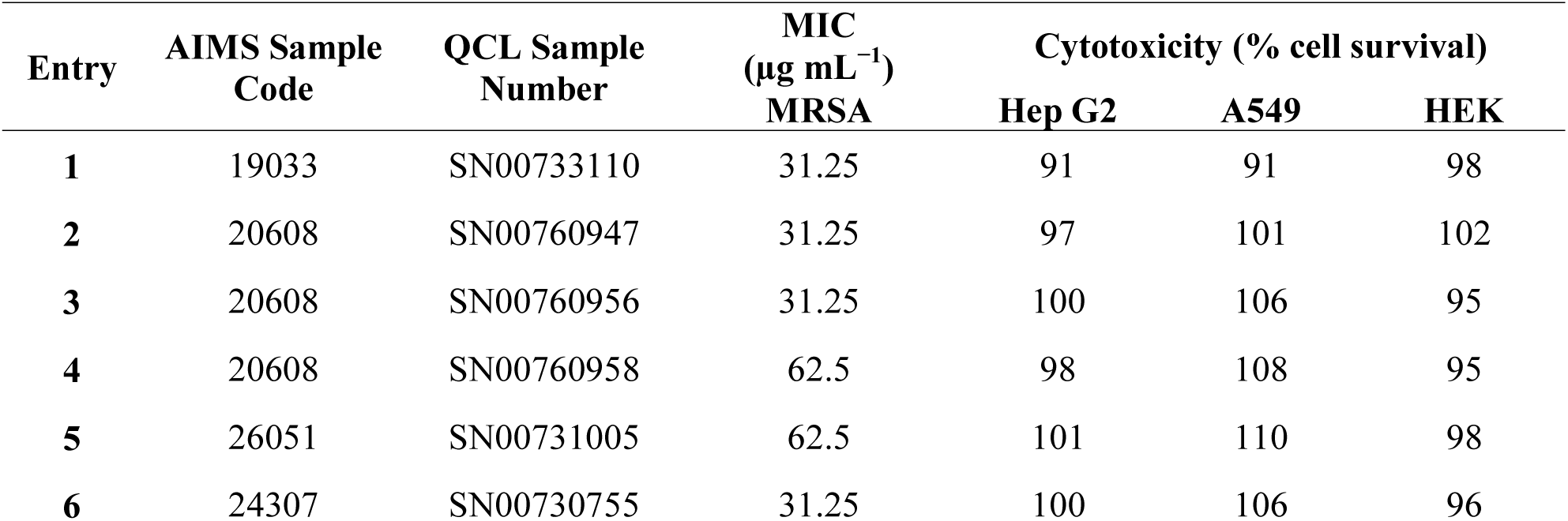
Summary of the Nine Marine Samples Selected for Further Study.

### 2.2. Isolation and Characterization of Bioactive Compounds

Following the primary screening of the AIMS library and selection of positive hits, HPLC was used to separate and isolate active compounds, guided by bioassays against MRSA. Extracts were fractionated by analytical HPLC (see Experimental section and Supporting Information for further details), and fractions evaluated for bioactivity. Preparative scale HPLC was carried out on each bulk sample to isolate the active component (Table S4, Figures S2– S7), and tandem mass spectrometry (MS/MS) methods used to deduce structures (Table 2 and Supporting Information).^19–20 21^ Insufficient quantities were obtained for NMR analyses.

Active components were isolated and characterised for five of the six extracts shown in Table 1: aromatic amines **1**/**2** from the *Lendenfeldia sp*. samples (AIMS Sample Code 20608, Table 1 entries 2-4); tertiary aliphatic amine **3** from the *Dysidea herbacea* extract (AIMS Sample Code 19033, Table 1 entry 1); and aliphatic tertiary amine **4**/ quaternary amine salt **5** from *Ircinia gigantea* (AIMS Sample Code 26051, Table 1 entry 5). The active component of the other active Demospongiae extract (AIMS Sample Code 24307, Table 1 entry 6) could not be isolated.

**Table 2.**
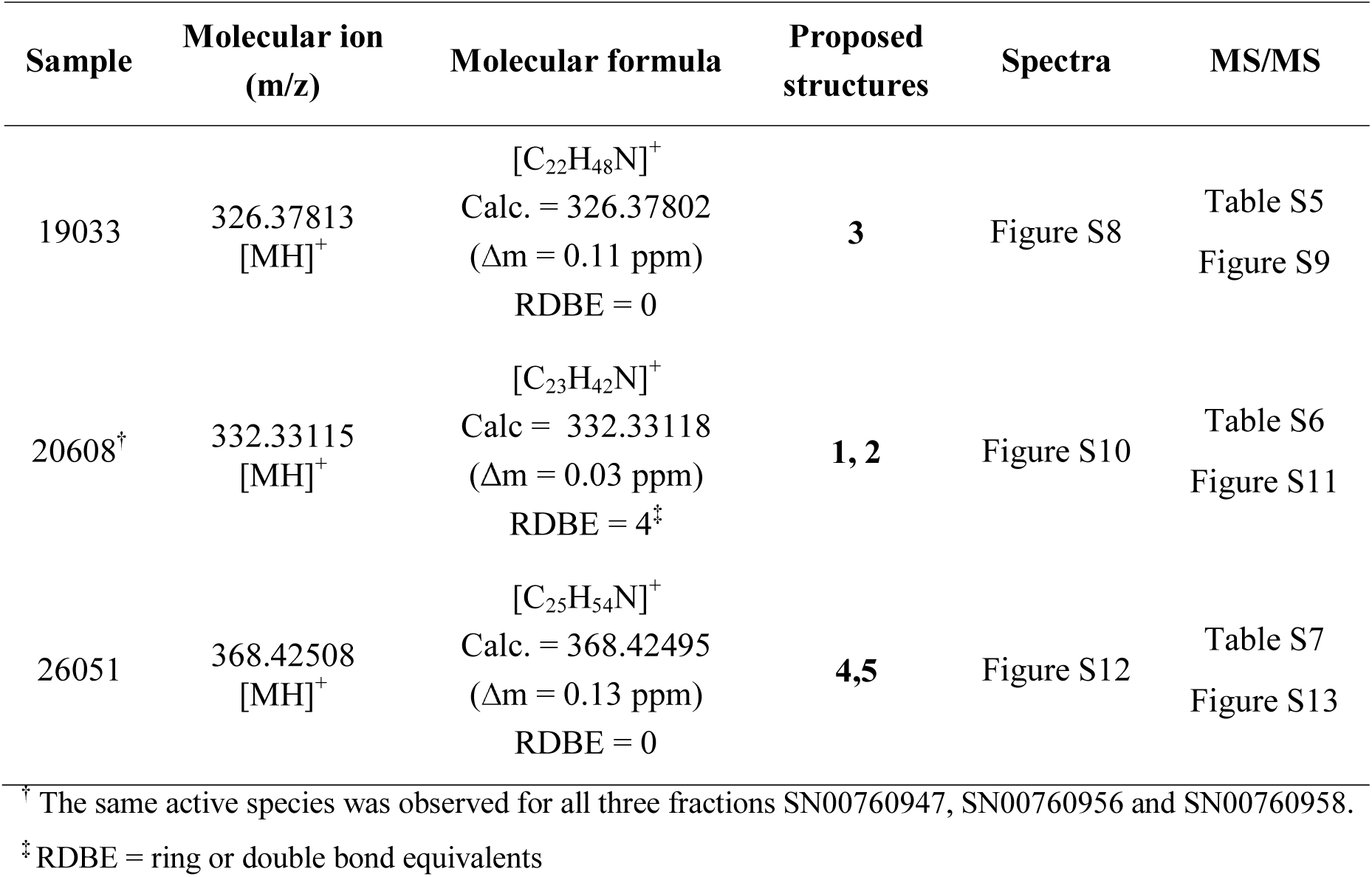
Key MS Data for Bioactive Samples, and Proposed Structures as Shown in Figure 1.

| Sample | Molecular ion<br>(m/z) | Molecular formula | Proposed<br>structures | Spectra | MS/MS |
| --- | --- | --- | --- | --- | --- |
| 19033 | 326.37813<br>[MH]<sup>+</sup> | [C<sub>22</sub>H<sub>48</sub>N]<sup>+</sup><br>Calc. = 326.37802<br>(Δm = 0.11 ppm)<br>RDBE = 0 | **3** | Figure S8 | Table S5<br>Figure S9 |
| 20608<sup>†</sup> | 332.33115<br>[MH]<sup>+</sup> | [C<sub>23</sub>H<sub>42</sub>N]<sup>+</sup><br>Calc = 332.33118<br>(Δm = 0.03 ppm)<br>RDBE = 4<sup>‡</sup> | **1, 2** | Figure S10 | Table S6<br>Figure S11 |
| 26051 | 368.42508<br>[MH]<sup>+</sup> | [C<sub>25</sub>H<sub>54</sub>N]<sup>+</sup><br>Calc. = 368.42495<br>(Δm = 0.13 ppm)<br>RDBE = 0 | **4,5** | Figure S12 | Table S7<br>Figure S13 |

<sup>†</sup> The same active species was observed for all three fractions SN00760947, SN00760956 and SN00760958.
<sup>‡</sup> RDBE = ring or double bond equivalents

### 2.3. Synthesis

To validate the structures proposed for the natural products, and to explore the potential of these a compounds as bioactive agents, a series of tertiary amine derivatives of compounds **1**–**5** were synthesised from 1-bromooctane **6**, *m*-toluidine **7**, *p*-toluidine **8**, benzyl amine **9**, 1-iododecane **10**, *N*,*N*-dihexylamine **11** and *N*,*N*,*N*-trioctylamine **12** (Schemes 1 and 2). *o*-Toluidine is carcinogenic and therefore was not used in synthetic experiments.

**Scheme 1.**
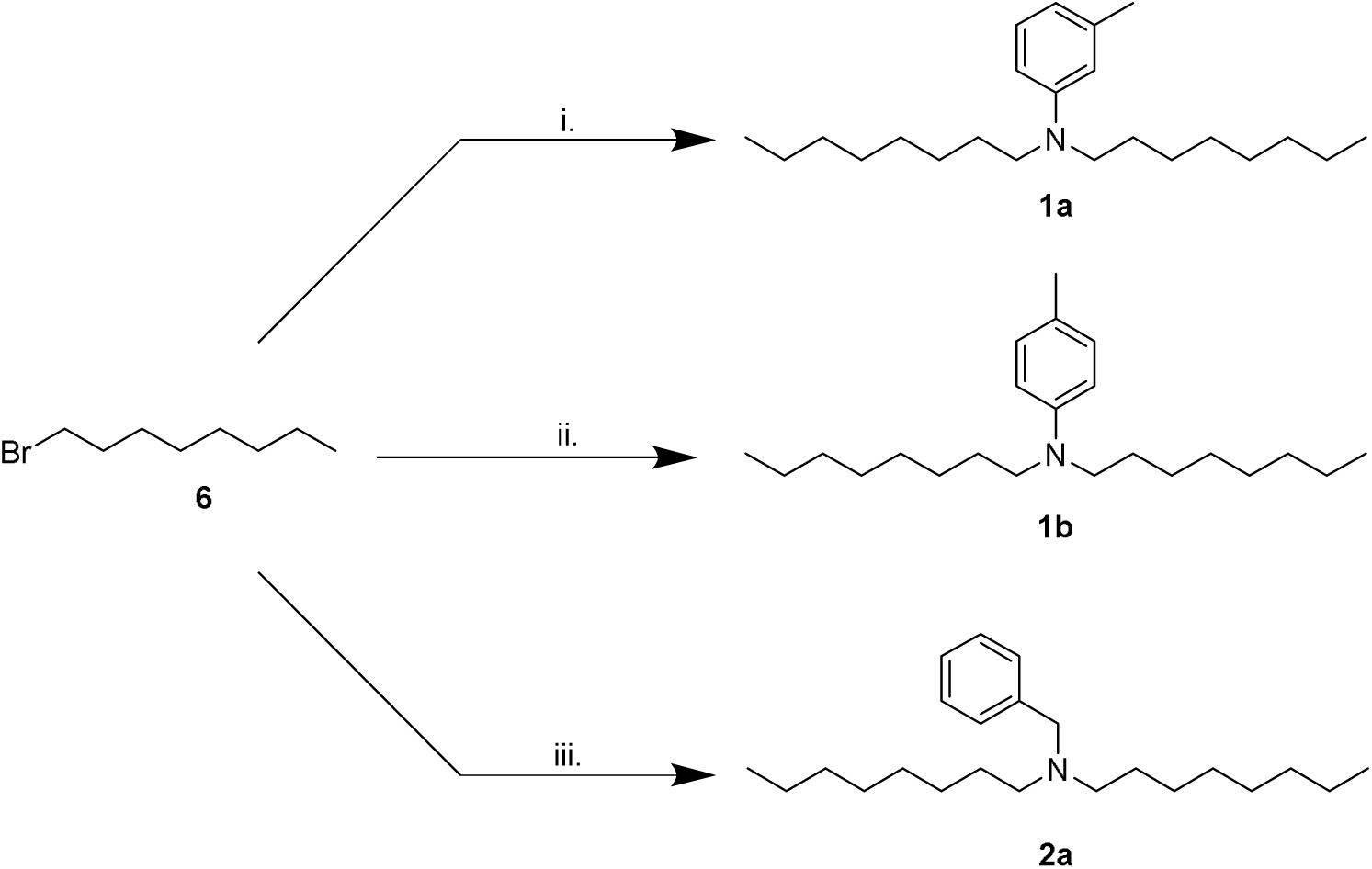
Synthesis of Amines Related to Natural Products 1 and 2. i. *m*-toluidine **7**, K_2_CO_3_, KI, MeCN, 60 °C, overnight, 8 %; ii. *p*-toluidine **8**, K_2_CO_3_, KI, MeCN, 60 °C, overnight, 6 %; iii. benzylamine **9**, K_2_CO_3_, MeCN, 82 °C, overnight, 19 %.

Compounds **1a**, **1b** and **2a** were prepared using 1-bromooctane **6** to alkylate *m*-toluidine **7**, *p*-toluidine **8**, benzyl amine **9** respectively (Scheme 1), giving three compounds based on the active component of AIMS sample 20608. Compound **3**, the active component of AIMS sample 19033, was prepared by reacting 1-iododecane **10** with *N,N*-dihexylamine **11** (Scheme 2), while *N,N,N*-trioctylamine hydrochloride **13** was prepared from the free amine **12** as a simple and readily accessible analogue of the natural product structures **4** and **5** that had been isolated from AIMS sample 26051.

**Scheme 2.**
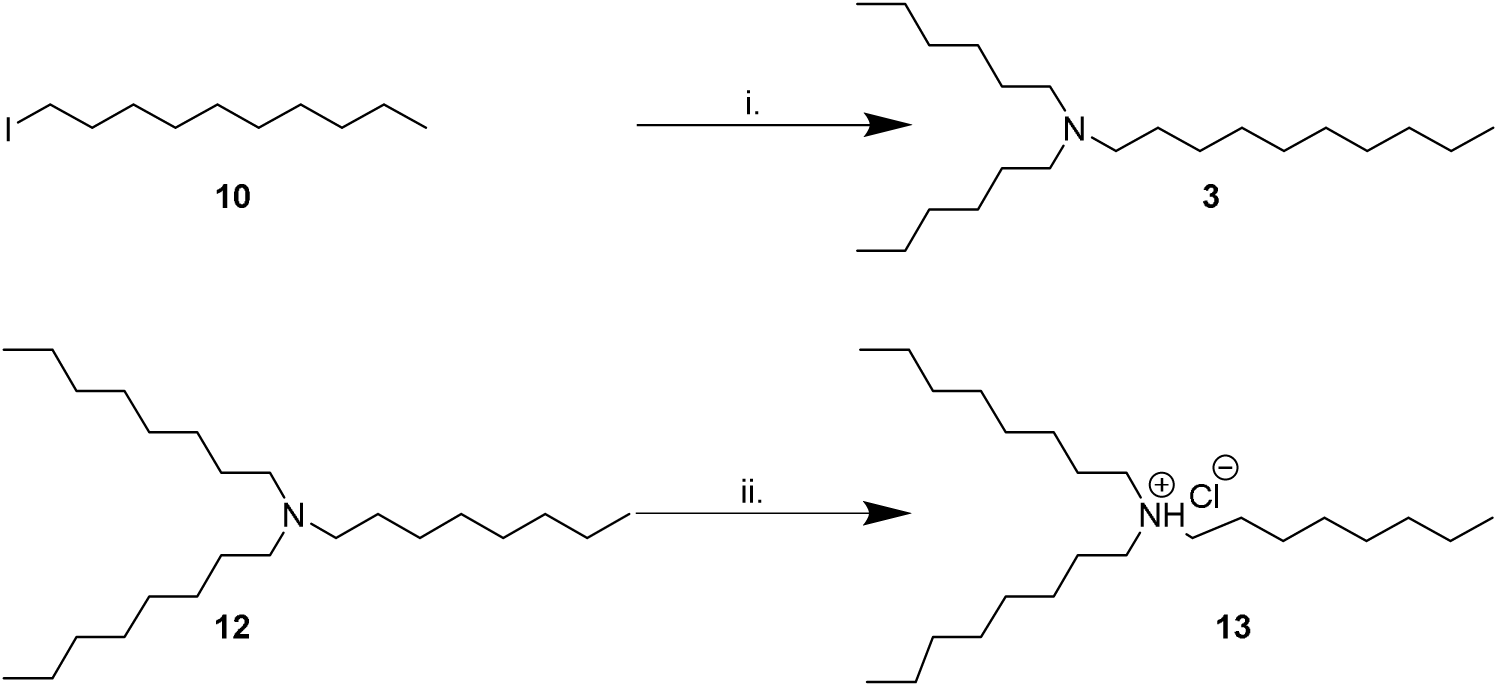
Synthesis of Amines Related to Natural Products 3–5. i. *N*,*N*-dihexylamine **11**, K_2_CO_3_, MeCN, 82 °C, overnight, 21 %; ii. HCl in 1,4-dioxane, 5 min, 99%.

Finally, to broaden the scope of this work, we sought to combine the tertiary amine structures elucidated in this study with a triazolyl naphthalimide pendant, recently shown to be an important component of a new class of anti-tubercular agents.^23–24^ Thus *N,N*-dioctylamine **14** was alkylated with 4-bromo-1-butyne **15**, and the resulting alkyne product **16** ‘clicked’^25^ with 6-azido-2-ethyl-1*H*-benzo[*de*]isoquinoline-1,3(2*H*)-dione **17**^26^ to afford the naphthalimide derivative **18** (Scheme 3).

**Scheme 3.**
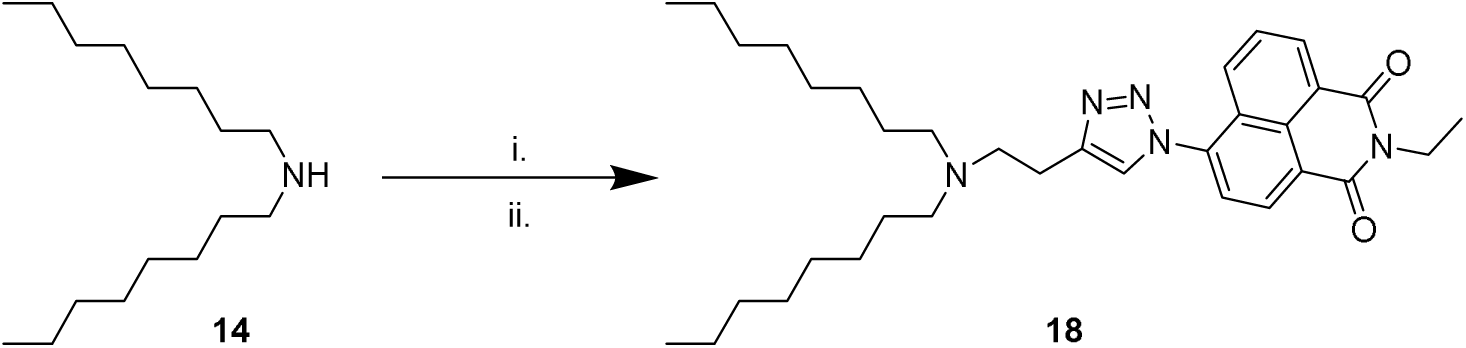
Synthesis of Naphthalimide Derivative 18. i. 4-bromo-1-butyne **15**, K_2_CO_3_, KI, MeCN, 60 °C, overnight, 2 %; ii. 6-azido-2-ethyl-1*H*-benzo[*de*]isoquinoline-1,3(2*H*)-dione **17**, CuSO_4_, sodium ascorbate, *^t^*BuOH/ H_2_O, 60 °C, overnight, 1 %.

While the yields of many synthetic steps were low, sufficient quantities of material were nonetheless isolated to enable characterisation, and biological evaluation, so the synthetic reactions were not further optimised.

### 2.4. Structural Comparison of Synthetic Compounds to Natural Products

Synthetic compounds were investigated using mass spectrometry (MS/MS and accurate mass) and analysis of biological activity. Comparing the major ions in the mass spectra of synthetic **1a** and **1b** (Table S8), **2a** (Table S9), **3** (Table S10) and **13** (Figure S11) with the natural products shows good correlation (Tables S6, S5 and S7, and Figures S9, S11 and S13 respectively). Some minor differences are apparent, which most likely arise due to differences in the amounts of material analysed (which are significantly greater for the synthesized products), and differences in the instrumentation used.

### 2.5. Antibacterial Activity and Toxicity of Synthetic Compounds

Synthetic compounds were assessed for bioactivity against MRSA, *P. aeruginosa*, uropathogenic *E. coli* and *M. tuberculosis*. Interestingly, all synthetic derivatives compounds showed similar MICs against MRSA (the organism against which the original natural product screening assays had been conducted), typically around 12.5 μM. The simple amine salt **13** proved the most effective of the synthetic compounds against *M. tuberculosis* with an MIC of 0.02 µM, and showed moderate inhibitory activity against the other bacteria. Compound **3** displayed broad activity, with low MICs against *P. aeruginosa* (MIC 3.1 µM), E. coli (6.2 µM) and *M. tuberculosis* (3.1 µM). The naphthalimide derivative **18** displayed good selective activity against *E. coli* (MIC 1.5 µM).

**Table 3.** Anti-Bacterial Activity of Synthetic Compounds Against MRSA, *P. Aeruginosa, E. Coli* and *M. Tuberculosis*.

| Synthetic compound | MIC ($\mu\text{M}$)<sup>†‡</sup> | | | |
| --- | --- | --- | --- | --- |
|  | MRSA | *M. tuberculosis* | *E. coli* | *P. aeruginosa* |
| **1a** | 12.5 | 1.5 | 12.5 | 6.2 |
| **1b** | 100 | 25 | 100 | 100 |
| **2a** | 12.5 | 1.5 | 50 | 50 |
| **3** | 12.5 | 3.1 | 6.2 | 3.1 |
| **13** | 12.5 | 0.02 | 25 | 25 |
| **18** | 12.5 | 12.5 | 1.5 | 12.5 |

<sup>†</sup> Samples were diluted from 100 $\mu\text{M}$ to 0.0002 $\mu\text{M}$ and incubated with bacteria (starting concentration OD<sub>600</sub>=0.001) under optimum assay conditions as described in the Experimental section.
<sup>‡</sup> Values represents averages from three independent repeats.

The potential toxicity of the synthetic compounds was also evaluated, against A549, THP-1, HepG2 and HEK 293 and cell lines (Table 3). None of the synthetic compounds showed significant toxicity against HepG2 or A549 cells. Synthetic compounds **1a**, **2a**, **3**, **13** and **18** all showed some toxicity against THP1 and/or HEK 293 cells, with Minimum Toxicity Concentrations (MTC) as low as 3.1 µM. Compound **1b** showed low toxicity against all four of these cell lines, but also low activity (Table 4). Compound **1a** showed only mild effects on all cell lines tested (MTC 50–100 µM), while also displayed a broad antibacterial profile, suggesting this compound may be a candidate for further investigation.

**Table 4.** Toxicity of Synthetic Compounds to Cell Lines.

| Synthetic samples | Minimum Toxicity Concentration of Drug (MTC $\mu\text{M}$)<sup>†‡§</sup> | | | |
| --- | --- | --- | --- | --- |
|  | A549 | THP1 | HepG2 | HEK 293 |
| **1a** | > 100 | 50 | > 100 | 50 |
| **1b** | > 100 | >100 | > 100 | > 100 |
| **2a** | > 100 | 6.5 | > 100 | 50 |
| **3** | >100 | 50 | > 100 | 25 |
| **13** | 50 | 3.1 | > 100 | 6.5 |
| **18** | > 100 | > 100 | > 100 | 3.1 |

<sup>†</sup> Cells seeded at a concentration of $2 \times 10^5$ cells/ well; samples diluted 100 $\mu\text{M}$ to 0.0002 $\mu\text{M}$ .
<sup>‡</sup> Cells incubated with sample under humidified incubation of 37 °C with 5 % $\text{CO}_2$ .
<sup>§</sup> Values represents averages of three independent repeats.

Comparing data from the antimicrobial activity and cytotoxicity assays shows that the active concentration ranges for these synthetic compounds against bacteria are substantially lower than active concentration ranges against the mammalian cells tested, particularly HepG2 and A549 cells. THP1 cells and the primary cells appeared more sensitive to these compounds. Cross-referencing the biological activity and toxicity data for these compounds suggest that they have some potential for further development.

Compounds **1a**, **13** and **18**, which showed the most promising activity against *M. tuberculosis*, were assessed against single and multi-drug resistant *M. tuberculosis* strains (Table 5). Compound **13** exhibited strong inhibitory activity against all resistant strains with MICs as low as 0.24 µm; Compound **1a** showed moderate and variable activity against all assay strains (MIC 6.7–20.0 µm) and **18** showed similar activity against all resistant strains with an MIC of 2.2 µm.

**Table 5.** Activity of synthetic compounds against drug resistant *M. tuberculosis* strains.

| *M. tb* strain<sup>†</sup> | Minimum inhibitory concentration (MIC) Drug ($\mu\text{M}$) | | |
| --- | --- | --- | --- |
|  | **1a** | **13** | **18** |
| sensitive | 20 | 0.74 | 2.2 |
| Inh | 20 | 0.74 | 6.7 |
| Rif, Inh | 20 | 0.74 | 6.7 |
| Rif, Inh, Eth | 20 | 0.74 | 6.7 |

<sup>†</sup> Rif = resistant to rifampicin; Inh = resistant to isoniazid; Eth = resistant to ethambutol;

## 3. CONCLUSIONS

Commercial drugs vancomycin and rifampin remain the reference standard for the treatment of invasive MRSA and *M. tuberculosis* infections respectively. Yet the number of vancomycin-resistant *S. aureus* (VRSA) and rifampicin-resistant *M. tuberculosis* strains is on the rise. The emergence of antibiotic resistance brings a need for novel, effective antibacterial agents that are resistant to antimicrobial resistance.

Thus we assayed 1434 extracts from the AIMS Bioresources Library^16^ against MRSA, finding three samples that have a promising combination of high antibacterial activity and low toxicity to mammalian cells: AIMS Sample Codes 20608, 26051 and 19033 (Table 1). Samples of these extracts were subjected to HPLC purification and bioassay guided fractionation, enabling bioactive components to be isolated in low yield («1 mg). Then high resolution MS and tandem MS analysis was used to decipher structures (Table 2, Figure 1). The proposed structures are all tertiary amines or quaternary amine salts: aromatic amines **1**/**2** (from *Lendenfeldia* sample number 20608), aliphatic amine **3** (from *Dysidea herbacea* sample number 19033), the aliphatic tertiary amine **4** and quaternary amine salt **5** (from *Ircinia sp*. sample number 26051).

Synthetic compounds based on the natural product structures **1**–**5** was prepared to validate and expand these findings. Synthetic compounds **1a**, **1b**, **2a**, **3** and **13** showed tandem MS fragmentation patterns consistent with the natural product samples, thus supporting the structures proposed for those samples, and promising bioactivity/ toxicity profiles. Naphthalimide derivative **18** was prepared as a hybrid of the amines uncovered in this study, and naphthalimide-amine derivatives we have previously reported as potent anti-mycobacterial agents.^23–24^

The compounds uncovered in this study add to the growing arsenal of antimicrobial agents from the sea,^2–3^ and offer interesting new avenues for further investigation in the quest for new, effective agents to combat the growing scourge of multidrug resistant bacteria.

## 4. EXPERIMENTAL METHODS

### 4.1. General

Chemical reagents were purchased from BDH Chemicals and Sigma Aldrich (Castle Hill, Sydney, Australia) and used as supplied unless otherwise indicated.

### 4.2. Natural Product Library

Natural product extracts were provided by the Australian Institute of Marine Science (AIMS), Townsville, Queensland as part of the AIMS Bioresources Library,^16^ via the Queensland Compound Library,^17^ (now called Compounds Australia^18^). Crude extracts had been partially fractionated by AIMS/ QCL to generate a library of 1434 samples, supplied in DMSO (100%) solution and stored at −80 °C. Original concentrations as provided were 5 mg mL^−1^. Stock solutions were made by diluting these samples by a factor of 1:10 in dH_2_O and stored at −80 °C.

### 4.3. Screening the AIMS Extract Library Against MRSA

Each test sample (10 µL) was dispensed into a separate well of a 96 well microtiter plates (final sample concentration 0.5 mg mL^−1^) using sterile dH_2_O. Bacterial suspension (90 µL, OD600nm 0.001) was added to each well and plates were incubated at 37 ^o^C for 18 hours. To determine MIC of samples, crude extracts were added to wells in sequential 2-fold dilutions and incubated with diluted bacteria as described previously.^23–24^ Resazurin (10 μL; 0.05% w/v) was added and plates were incubated for 3 h at 37 °C. The bioactivity of extracts was calculated by visual determination of colour change within wells or detection of fluorescence at 590 nm using a FLUOstar Omega microplate reader (BMG Labtech, Germany).

### 4.4. Evaluating Toxicity of AIMS Extract Library

Human alveolar epithelial cells (A549), ^27^ Madin-Darby canine kidney epithelial cells (MDCK),^28^ human leukaemia cells (THP-1),^29^ human hepatocellular carcinoma cells (Hep-G2),^30^ and human embryonic kidney cells 293 (HEK293)^31^ were grown and differentiated in complete RPMI (Roswell Park Memorial Institute Medium) and DMEM (Dulbecco's Modified Eagle's medium) tissue culture media (RPMIc and DMEMc). To determine toxicity of the AIMS extract library, 2 × 10^5^ of each cell type were added to a 96-well plate and left for 48 h at 37 °C to adhere. Extract samples at a final concentration of 0.5 mg mL^-1^ were added to the wells, then incubated for 7 days in a humidified 5% CO2 incubator at 37 °C. Then resazurin (10 μL of 0.05% w/v) was added and after 4 h, fluorescence measured as described previously. Cell viability was calculated as percentage fluorescence relative to untreated cells.

### 4.4. Purification of Natural Products from Extracts and Structure Elucidation

#### 4.4.1. High Performance Liquid Chromatography (HPLC) Purification

Samples were separated using analytical (Waters 2695 Alliance with Waters 2996 PDA, Sunfire reversed-phase column, and WFIII fraction collector) and preparative (Waters 600 HPLC pump, Phenomenex reversed-phase column, Waters 2487 UV detector and WFIII fraction collector) HPLC systems with UV detectors at 254 and 280 nm, employing a gradient of solvents A (dH_2_O) and B (acetonitrile) with trifluoroacetic acid (0.01%). Extract mixtures were kept at 4 °C until injection, then extract sample (100 μL) was injected onto an analytical Waters X-bridge C18 100 A (4.6 × 250 mm, 5 µm) reversed-phase column on the same analytical HPLC system described above. The mobile phase was obtained using 100% acetonitrile and 0% water at a flow rate of 1 ml min^-1^ at 30 ºC over 80 min. Fractions, separated every 60 s, were collected. Purified fractions were flash-frozen in liquid nitrogen then freeze-dried overnight. The resulting fractionated extracts were re-suspended in DMSO and antibacterial activity versus MRSA was determined as described above.

Fractions identified as active against MRSA were further purified on the preparative HPLC unit described above, using a C18 100 A (250 × 21.2 mm, 10 µm) reversed-phase column (Phenomenex) with UV detection at 254 and 280 nm, 7 mL min^-1^ flow rate with water/acetonitrile gradient containing 0.1% trifluoroacetic acid.

The gradient for AIMS extracts 19033, 20608, and 26051 was 0% B initially, increased to 40% B over 20 min, then to 100% at 60 min, held at 100% for 10 min, and finally a linear decrease to 0% B over 5 min and held at 0% B for 10 min prior to the next run. Compounds thus purified were evaluated for biological activity and analysed by MS to determine potential structures for the bioactive components.

#### 4.4.2. Identification and Structure Elucidation

Purified compounds were identified and characterised using MS. High resolution ESI mass spectra (HRMS) were recorded on a Bruker Apex Qe 7T Fourier Transform ion cyclotron resonance mass spectrometer with an Apollo II ESI MTP ion source with samples (in CH3CN:H_2_O 1:1) infused using a Cole Palmer syringe pump at 180 µL h^-1^. Where required, low resolution ESI tandem MS was performed on a Bruker amaZon SL ion trap via syringe infusion or by injection into a constant flow stream with a rheodyne valve and an Alltech HPLC pump (mobile phase methanol, flow rate 0.3 mL min^-1^) connected to an Apollo II ESI MTP ion source in positive ion mode. Tandem mass spectra of the [M+H]^+^ parent ion were obtained manually up to MS^5^ (depending on sensitivity). Spectra were acquired in positive ion mode using a 1–4 Da isolation window, with the excitation amplitude manually optimized for each spectrum to have the selected mass at ~10% of the height of the largest fragment. Data analysis was performed for both high resolution MS and low resolution tandem MS data using Bruker DataAnalysis 4.0 with smart formula assuming C, H, N, O, Na (0-1), mass error <2 ppm, C:H ratio 3 maximum, even electron (or both for tandem MS data). The results of high resolution MS data analysis were further refined manually by comparing isotopic fine structures of simulations where possible (resolving power > 200,000) to further eliminate potential formulae within the 2ppm mass error window (particularly ^15^N, ^18^O, ^2^H, ^13^C and ^13^C_2_ isotopes and confirm no ^34^S presence).

### 4.5. Synthesis

#### 4.5.1. *N,N*-Dioctyl-3-methylaniline 1a

To a solution of 1-bromooctane 6 (6.3 mL, 36.6 mmol) in acetonitrile (50 mL) was added potassium carbonate (25.3 g, 183 mmol), KI (6.08 g, 36.6 mmol) and m-toluidine 7 (1.96 mL, 18.3mmol) stirred at 60 °C overnight. The suspension was filtered and washed with acetonitrile (3 × 50 mL), then concentrated by rotary evaporation to yield the crude product (1.50 g). Purification by automated column chromatography (100 g cartridge, 100% petroleum benzine over 12 CV) then by preparative TLC (10% ethyl acetate:petroleum benzine) afforded the pure compound 1a as a yellow oil (0.46 g, 8 %). ^1^H NMR (500 MHz, CDCl_3_): δ 7.13 – 7.07 (m, 1 H), 6.50 – 6.44 (m, 3 H), 3.28 – 3.22 (m, 4 H), 2.32 (s, 3 H), 1.64 – 1.53 (m, 4 H), 1.38 – 1.24 (m, 20 H), 0.94 – 0.87 (m, 6 H); ^13^C NMR (125 MHz, CDCl_3_): δ 148.3, 138.7, 129.0, 116.0, 112.4, 109.0, 51.0, 31.8, 29.5, 29.3, 27.3, 27.2, 22.7, 22.0, 14.1; LRMS (ESI+): m/z 332.33 [M+H]^+^, 100%; HRMS (ESI): m/z calculated for [C_23_H_42_N]^+^, [M + H]^+^ 332.3317, found 332.33054.

#### 4.5.2. *N,N*-Dioctyl-4-methylaniline 1b

To a solution of 1-bromooctane 6 (6.3 mL, 36.6 mmol) in acetonitrile (50 mL) was added potassium carbonate (25.3 g, 183 mmol), KI (6.08 g, 36.6 mmol) and p-toluidine 8 (1.96g, 18.3 mmol) stirred at 60 °C overnight. The suspension was filtered then concentrated by rotary evaporation to yield the crude product (1.65 g). Purification by automated column chromatography (100 g cartridge, 100% petroleum benzine over 4 CV, ramping to 100% ethyl acetate over 4 CV) gave the product 1b as a yellow oil (0.39 g, 6 %). ^1^H NMR (500 MHz, CDCl_3_): δ 7.02 (d, J = 8.2 Hz, 2 H), 6.68 – 6.47 (m, 2 H), 3.33 – 3.10 (m, 4 H), 2.25 (s, 3 H), 1.58 – 1.52 (m, 4 H), 1.40 – 1.20 (m, 20 H), 0.95 – 0.82 (m, 6 H); ^13^C NMR (125 MHz, CDCl_3_): δ 146.2, 129.7, 124.3, 112.2, 51.3, 31.9, 29.5, 29.4, 27.3, 27.2, 22.7, 20.1, 14.1; LRMS (ESI+): m/z 332.33 [M+H]^+^, 100%; HRMS (ESI): m/z calculated for [C_23_H_42_N]^+^ [M+H]^+^ 332.3317, found 332.33095.

#### 4.5.3. *N*-Benzyl-*N*-octyloctan-1-amine 2a

To a solution of 1-bromooctane 6 (6.3 mL, 36.6 mmol) in acetonitrile (50 mL) were added potassium carbonate (25.3 g, 183 mmol) and benzylamine 9 (2 mL, 18 mmol) then stirred at 60 ºC overnight. The mixture, a clear colourless solution, was concentrated using the rotary evaporator, to give a white solid. The solid was triturated with DCM (1 × 50 mL, 1 × 25 mL) and the DCM solution concentrated on the rotary evaporator to yield yellow oil (5.1 g). Purification by automated column chromatography (100 g cartridge, 0–40% ethyl acetate (EtOAc) in petroleum benzine over 10 CV) yielded N-benzyl-N-octyloctan-1-amine 2a as a colourless oil (1.15 g, 19 %). ^1^H NMR (500 MHz, CDCl_3_): δ 7.36 – 7.29 (m, 4H), 7.26 – 7.21 (m, 1H), 3.56 (s, 2H), 2.47 – 2.35 (m, 4H), 1.54 – 1.41 (m, 4H), 1.36 – 1.21 (m, 20H), 0.90 (t, J = 7.0 Hz, 6H); ^13^C NMR (125 MHz, CDCl_3_): δ 140.3, 128.8, 128.0, 126.6, 58.6, 53.8, 31.9, 29.6, 29.3, 27.5, 27.0, 22.7, 14.1; LRMS (ESI+): m/z 332.33 [M+H]^+^, 100%; HRMS (ESI): m/z calculated for [C_23_H_42_N]^+^ [M + H]^+^ 332.3317, found 332.3306.

#### 4.5.4. *N,N*-Dihexyldecan-1-amine 3

To a solution of 1-iododecane 10 (2.75 mL, 12.9 mmol) in acetonitrile (60 mL) was added potassium carbonate (16.5 g, 129 mmol) and dihexylamine 11 (3 mL, 12.9 mmol), then the mixture was stirred at reflux overnight. The suspension was filtered to remove K_2_CO_3_ and washed with acetonitrile (3 × 50 mL), then concentrated by rotary evaporation to yield the crude product (7.8 g). The crude product was purified by automated column chromatography (100 g cartridge, 100% petroleum benzene 2CV, then 0–60% ethyl acetate in petroleum benzene over 10 CV) to yield the product as an oil (0.89 g, 21 %). ^1^H NMR (500 MHz, CDCl_3_): δ 2.41 – 2.37 (m, 6 H), 1.38– 1.46 (m, 6 H), 1.35 – 1.21 (m, 28 H), 0.91 – 0.86 (m, 9 H); ^13^C NMR (125 MHz, CDCl_3_): δ 54.2,54.2, 31.9, 31.9, 29.7, 29.6, 29.6, 29.3, 27.7, 27.4, 26.9, 22.7, 14.1, 14.1; LRMS (ESI+): m/z 326.41 [M+H]^+^, 100%; HRMS (ESI): m/z calculated for [C_22_H_48_N]^+^ [M + H]^+^ 326.3787, found 326.3776.

#### 4.5.5. *N,N,N-*Trioctylammonium chloride 13

To a solution of *N,N,N*-trioctylamine 12 (1.0 g, 2.80 mmol) in 1,4-dioxane (1.0 mL) was added 4M HCl in dioxane (2.80 mL, 11.2 mmol) in an ice bath. Instantaneously a precipitate formed and after 5 min this was collected by vacuum filtration to yield a white solid (1.08 g, 99 % yield). ^1^H NMR (500 MHz, CDCl_3_) δ 11.40 (br s, 1 H), 2.90 (td, J = 4.7, 12.5 Hz, 6 H), 1.74 – 1.66 (m, 6 H), 1.29 – 1.13 (m, 30 H), 0.79 (t, J = 7.0 Hz, 9 H); ^13^C NMR (125 MHz, CDCl_3_) δ 52.2, 31.4, 28.8, 28.7, 26.6, 23.0, 22.3, 13.8.

#### 4.5.6. *N*-(But-3-yn-1-yl)-*N*-octyloctan-1-amine 16

To a flask charged with potassium carbonate (4.4 g, 31.8 mmol) and potassium iodide (880 mg, 5.30 mmol) was added acetonitrile (72 mL) followed by dioctylamine 14 (8 mL, 26.5 mmol) and 4-bromo-1-butyne 15 (2.74 mL, 29.2 mmol). The suspension was stirred at 60°C for 18 h, then filtered and washed with acetonitrile (3 × 50 mL), and concentrated by rotary evaporation to yield the crude product. This was purified by automated column chromatography (100 g cartridge, 0%–15% ethyl acetate in petroleum benzine over 10 CV) to yield the product as a yellow oil (0.15 g, 2%). ^1^H NMR (500 MHz, CDCl_3_): δ 2.64 – 2.57 (m, 2 H), 2.39 – 2.31 (m, 4 H), 2.23 (dt, J = 2.7, 7.6 Hz, 2 H), 1.88 (t, J = 2.6 Hz, 1 H), 1.41 - 1.31 (m, 4 H), 1.27 – 1.14 (m, 20 H), 0.81 (t, J = 7.0 Hz, 6 H); ^13^C NMR (125 MHz, CDCl_3_): δ 83.3, 68.7, 54.0, 52.7, 31.8, 29.6, 29.3, 27.6, 27.2, 22.6, 16.7, 14.1; LRMS (ESI+): m/z 294.29 [M+H]^+^, 100%; HRMS (ESI): m/z calculated for C_20_H_40_N^+^ [MH]^+^ 294.3161, found 294.3157.

#### 4.5.7. 6-(4-(2-(Dioctylamino) ethyl)-1*H*-1,2,3-triazol-1-yl)-2-ethyl-1*H*-benzo[de]isoquinoline-1,3(*2H*)-dione 18

To a solution of 16 (400 mg, 1.24 mmol) and 6-azido-2-ethyl-1*H*-benzo[de]isoquinoline-1,3(*2H*)-dione 17 (365 mg, 1.61 mmol) in tert-butanol: water (12.4 mL) were added copper sulphate hexahydrate (31.0 mg, 0.12 mmol) and ascorbic acid sodium salt (73.8 mg, 0.37 mmol) then the solution was stirred at 60ºC overnight. Precipitants were removed by filtration and washed with acetonitrile (3 × 50 mL), then the filtrate was concentrated by rotary evaporation and purified by automated column chromotography (100g cartridge, 0%–20% methanol in dichloromethane over 8 CV). Purified fractions were re-purified by automated reversed phase chromotography (30g cartridge C18 silica (Biotage), 0%–90% acetonitrile in water over 17 CV) and concentrated to yield the product as a yellow solid (10 mg, 1%). ^1^H NMR (500 MHz, CDCl_3_): δ 8.73 (d, J = 7.6 Hz, 2 H), 8.21 (dd, J = 0.9, 8.5 Hz, 1 H), 8.09 (s, 1 H), 7.89 – 7.81 (m, 2 H), 4.29 (q, J = 7.1 Hz, 2 H), 3.67 - 3.56 (m, 2 H), 3.49 3.37 (m, 2 H), 3.25 - 3.06 (m, 4 H), 1.87 - 1.70 (m, 4 H), 1.38 (t, J = 7.0 Hz, 3 H), 1.34 – 1.23 (m, 10 H), 0.89 (t, J = 7.0 Hz, 6 H); ^13^C NMR (126 MHz, CDCl_3_): δ 162.4, 161.9, 142.1, 136.8, 131.2, 129.6, 128.1 (128.09), 128.1 (128.08), 127.7, 125.4, 124.1, 123.2, 122.6, 122.1, 51.6, 51.3, 34.8, 30.6, 28.7, 28.0, 25.7, 22.0, 21.5, 20.1, 13.0, 12.3; LRMS (ESI+): m/z 560.38 [M+H]^+^, 100%; HRMS (ESI): m/z calculated for C_34_H_50_N_5_O_2_^+^ [MH]^+^ 560.3965, found 560.3950.

### 4.6. Screening of Synthetic Compounds

MRSA (provided by Dr John Merlino at Concord Hospital, Sydney), *P. aeruginosa* PAO1 (provided by Dr Jim Manos, University of Sydney) and *E. coli* EC958 (provided by Professor Mark Schembri, University of Queensland) were grown in LB media. *M. tuberculosis* H37Rv was grown in Middlebrook 7H9 media (Bacto, Australia) containing albumin, dextrose, and catalase (ADC), 20% Tween 80, and 50% glycerol (Sigma-Aldrich, Australia). The synthetic samples were suspended in DMSO and diluted in series to final concentrations of 100 µM – 0.001µM, using sterile dH_2_O. Antibacterial activities and toxicity were determined for each compound via broth dilution resazurin assay as described above.

## Supporting information

## ASSOCIATED CONTENT

### Supporting Information

Bioactivity and toxicity screening data, HPLC fractionation and purification protocols, plus mass spectrometry data (HRMS spectra, tables of daughter ions, and proposed fragmentation pathways) for natural products and synthetic compounds.

## AUTHOR INFORMATION

Corresponding Author

PJR: phone +61 2 9351 5020;

JAT: phone +61 2 9036 6582

### Author Contribution Statement

PJR, JAT, MD, NP (mass spectrometry) and MPS (synthesis) conceived and designed the experiments; MD, NP (mass spectrometry), MPS (synthesis) and GT (bioassays with virulent *M. tuberculosis*) performed the experiments; MD, NP, MPS, GT, JAT and PJR analyzed the data; MD, PJR, MPS and JAT wrote the paper.

### Notes

The authors declare no competing financial interest

## ACKNOWLEDGEMENT

We thank Dr Cody Szczepina for his assistance developing HPLC methods for natural product isolation and purification. We thank Dr John Merlino (Concord Hospital, Sydney), Dr Jim Manos (University of Sydney) and Professor Mark Schembri (University of Queensland) for provision of the MRSA*, P. aeruginosa* PAO1 and *E. coli* EC958 strains respectively. This work was supported by the National Health and Medical Research Council (NHMRC) Project APP1084266, the NHMRC Centre of Research Excellence in Tuberculosis Control (APP1043225), and the University of Sydney. MD was supported by an International Postgraduate Research Scholarship (IPRS) and Australian Postgraduate Award (APA) from the Australian Government.

## ABBREVIATIONS

A549: human alveolar epithelial cells
AIMS: Australian Institute for Marine Science
DMEM: Dulbecco's Modified Eagle's medium
HEK293: human embryonic kidney cells 293
HPLC: high-performance liquid chromatography
HTS: high-throughput screening
MDCK: Madin-Darby canine kidney epithelial cells
MIC: minimum inhibitory concentration
MRSA: methicillin resistant *Staphylococcus aureus*
MS: mass spectrometry
MS/MS: tandem mass spectrometry
MTC: minimum toxic concentration
NMR: nuclear magnetic resonance
RPMI: Roswell Park Memorial Institute Medium
THP-1: human leukaemia cells
Hep-G2: human hepatocellular carcinoma cells
VRSA: vancomycin-resistant *Staphylococcus aureus*

## REFERENCES

1. Molinski, T. F.; Dalisay, D. S.; Lievens, S. L.; Saludes, J. P., Drug development from marine natural products. Nat. Rev. Drug Discovery 2008, 8, 69–85.

2. Hughes, C. C.; Fenical, W., Antibacterials from the Sea. Chem. Eur. J. 2010, 16 (42), 12512–12525.

3. Indraningrat, A.; Smidt, H.; Sipkema, D., Bioprospecting Sponge-Associated Microbes for Antimicrobial Compounds. Mar. Drugs 2016, 14 (5), 87.

4. Mehbub, M. F.; Perkins, M. V.; Zhang, W.; Franco, C. M. M., New marine natural products from sponges (Porifera) of the order Dictyoceratida (2001 to 2012); a promising source for drug discovery, exploration and future prospects. Biotechnol. Adv. 2016, 34 (5), 473–491.

5. Schroeder, G.; Bates, S. S.; La Barre, S., Bioactive Marine Molecules and Derivatives with Biopharmaceutical Potential. In Blue Biotechnology, Barre, S. L.; Bates, S. S., Eds. 2018.

6. Newman, D. J.; Cragg, G. M., Natural Products as Sources of New Drugs from 1981 to 2014. J. Nat. Prod. 2016, 79 (3), 629–661.

7. Pye, C. R.; Bertin, M. J.; Lokey, R. S.; Gerwick, W. H.; Linington, R. G., Retrospective analysis of natural products provides insights for future discovery trends. Proc. Nat. Acad. Sci. U.S.A. 2017, 114 (22), 5601–5606.

8. Blunt, J. W.; Copp, B. R.; Keyzers, R. A.; Munro, M. H. G.; Prinsep, M. R., Marine natural products. Nat. Prod. Rep. 2017, 34 (3), 235–294.

9. Blunt, J. W.; Carroll, A. R.; Copp, B. R.; Davis, R. A.; Keyzers, R. A.; Prinsep, M. R., Marine natural products. Nat. Prod. Rep. 2018, 35 (1), 8–53.

10. McGivern, J. G., Ziconotide: a review of its pharmacology and use in the treatment of pain. Neuropsychiatr. Dis. Treat. 2007, 3 (1), 69–85.

11. Gordon, E. M.; Sankhala, K. K.; Chawla, N.; Chawla, S. P., Trabectedin for Soft Tissue Sarcoma: Current Status and Future Perspectives. Adv. Ther. 2016, 33 (7), 1055–1071.

12. Bassetti, M.; Baguneid, M.; Bouza, E.; Dryden, M.; Nathwani, D.; Wilcox, M., European perspective and update on the management of complicated skin and soft tissue infections due to methicillin-resistant Staphylococcus aureus after more than 10 years of experience with linezolid. Clin Microbiol Infect 2014, 20 Suppl 4, 3–18.

13. Dookie, N.; Rambaran, S.; Padayatchi, N.; Mahomed, S.; Naidoo, K., Evolution of drug resistance in Mycobacterium tuberculosis: a review on the molecular determinants of resistance and implications for personalized care. J. Antimicrob. Chemother. 2018, 73 (5), 1138–1151.

14. Pourakbari, B.; Mamishi, S.; Mohammadzadeh, M.; Mahmoudi, S., First-Line Anti-Tubercular Drug Resistance of Mycobacterium tuberculosis in IRAN: A Systematic Review. Front. Microbiol. 2016, 7, 1139.

15. Lopez-Avalos, G.; Gonzalez-Palomar, G.; Lopez-Rodriguez, M.; Vazquez-Chacon, C. A.; Mora-Aguilera, G.; Gonzalez-Barrios, J. A.; Villanueva-Arias, J. C.; Sandoval-Diaz, M.; Miranda-Hernández, U.; Alvarez-Maya, I., Genetic diversity of Mycobacterium tuberculosis and transmission associated with first-line drug resistance: a first analysis in Jalisco, Mexico. J. Glob. Antimicrob. Resist. 2017, 11, 90–97.

16. Evans-Illidge, E. A.; Logan, M.; Doyle, J.; Fromont, J.; Battershill, C. N.; Ericson, G.; Wolff, C. W.; Muirhead, A.; Kearns, P.; Abdo, D.; Kininmonth, S.; Llewellyn, L., Phylogeny drives large scale patterns in Australian marine bioactivity and provides a new chemical ecology rationale for future biodiscovery. PLoS One 2013, 8 (9), e73800.

17. Simpson, M.; Poulsen, S.-A., An Overview of Australia’s Compound Management Facility: The Queensland Compound Library. ACS Chem. Biol. 2014, 9 (1), 28–33.

18. Compounds Australia. https://www.griffith.edu.au/griffith-sciences/compounds-australia (accessed 20 September 2018).

19. Bouslimani, A.; Sanchez, L. M.; Garg, N.; Dorrestein, P. C., Mass spectrometry of natural products: current, emerging and future technologies. Nat. Prod. Rep. 2014, 31 (6), 718–729.

20. Kind, T.; Fiehn, O., Advances in structure elucidation of small molecules using mass spectrometry. Bioanal. Rev. 2010, 2 (1), 23–60.

21. Herbert Júnior, D.; Nathalya Isabel de, M.; Antônio Eduardo Miller, C., Electrospray Ionization Tandem Mass Spectrometry as a Tool for the Structural Elucidation and Dereplication of Natural Products: An Overview. In Tandem Mass Spectrometry - Applications and Principles, Prasain, J., Ed. InTech: 2012.

22. Kernan, M. R.; Faulkner, D. J., Sesterterpene sulfates from a sponge of the family Halichondriidae. J. Org. Chem. 1988, 53 (19), 4574–4578.

23. Yu, M.; Nagalingam, G.; Ellis, S.; Martinez, E.; Sintchenko, V.; Spain, M.; Rutledge, P. J.; Todd, M. H.; Triccas, J. A., Nontoxic Metal–Cyclam Complexes, a New Class of Compounds with Potency against Drug-Resistant Mycobacterium tuberculosis. J. Med. Chem. 2016, 59 (12), 5917–5921.

24. Spain, M.; Wong, J. K. H.; Nagalingam, G.; Batten, J. M.; Hortle, E.; Oehlers, S. H.; Jiang, X. F.; Murage, H. E.; Orford, J. T.; Crisologo, P.; Triccas, J. A.; Rutledge, P. J.; Todd, M. H., Antitubercular Bis-Substituted Cyclam Derivatives: Structure–Activity Relationships and in Vivo Studies. J. Med. Chem. 2018, 61 (8), 3595–3608.

25. Wong, J. K.-H.; Ast, S.; Yu, M.; Flehr, R.; Counsell, A. J.; Turner, P.; Crisologo, P.; Todd, M. H.; Rutledge, P. J., Synthesis and Evaluation of 1,8-Disubstituted-Cyclam/Naphthalimide Conjugates as Probes for Metal Ions. ChemistryOpen 2016, 5 (4), 375–385.

26. Yu, M.; Yu, Q.; Rutledge, P. J.; Todd, M. H., A Fluorescent “Allosteric Scorpionand” Complex Visualizes a Biological Recognition Event. ChemBioChem 2013, 14 (2), 224–229.

27. Giard, D. J.; Aaronson, S. A.; Todaro, G. J.; Arnstein, P.; Kersey, J. H.; Dosik, H.; Parks, W. P., In vitro cultivation of human tumors: establishment of cell lines derived from a series of solid tumors. J Natl Cancer Inst 1973, 51 (5), 1417–23.

28. Gaush, C. R.; Hard, W. L.; Smith, T. F., Characterization of an established line of canine kidney cells (MDCK). Proc Soc Exp Biol Med 1966, 122 (3), 931–5.

29. Tsuchiya, S.; Yamabe, M.; Yamaguchi, Y.; Kobayashi, Y.; Konno, T.; Tada, K., Establishment and characterization of a human acute monocytic leukemia cell line (THP-1). Int J Cancer 1980, 26 (2), 171–6.

30. Aden, D. P.; Fogel, A.; Plotkin, S.; Damjanov, I.; Knowles, B. B., Controlled synthesis of HBsAg in a differentiated human liver carcinoma-derived cell line. Nature 1979, 282 (5739), 615–6.

31. Graham, F. L.; Smiley, J.; Russell, W. C.; Nairn, R., Characteristics of a human cell line transformed by DNA from human adenovirus type 5. J Gen Virol 1977, 36 (1), 59–74.

