## Supplementary material for "Discovery of antimicrobial compounds from *Lendenfeldia*, *Ircinia* and *Dysidea* sponges using bioassay guided fractionation of marine extracts"

Mojdeh Dinarvand,^†,‡^ Nicholas Proschogo,^†^ Malcolm P. Spain,^†^ Gayathri Nagalingam,^‡,§^ Elena Martinez,^∥,¶^ Vitali Sintchenko,^∥,¶^ James A. Triccas^‡,§, ¶,*^ and Peter J. Rutledge^†,*^

^†^ School of Chemistry, The University of Sydney, Sydney, NSW 2006, Australia

^‡^ Department of Infectious Diseases and Immunology, Faculty of Medicine and Health, The University of Sydney, NSW, Australia

^§^ Tuberculosis Research Program, Centenary Institute, The University of Sydney, NSW, Australia

^∥^ Centre for Infectious Diseases and Microbiology Laboratory Services, Westmead Hospital, Westmead, NSW 2145, Australia

^¶^Marie Bashir Institute for Infectious Diseases and Biosecurity, The University of Sydney, NSW, Australia

**Contents**

1. Bioactivity and Toxicity Screening	2
2. HPLC Fractionation and Purification	6
3. Mass Spectrometry Data for Natural Products	10
4. Mass Spectrometry Data for Synthetic Compounds	18

### 1. BIOACTIVITY AND TOXICITY SCREENING.

**Table S1. Determination of MIC values for selected compounds against MRSA. MRSA was incubated with samples as described in the Experimental section.**

| Entry | AIMS Sample Code | Phylum | QCL Sample Number | Fraction | MRSA MIC  (µg mL^−1^) |
| --- | --- | --- | --- | --- | --- |
| 1 | 22565 | Porifera | SN00739718 | Crude Extract | 31.3 |
| 2 | 24606 | Porifera | SN00732793 | MeOH Eluent | 125.0 |
| 3 | 24307 | Porifera | SN00730771 | MeOH Eluent | 250.0 |
| 4^†^ | 20608 | Porifera | SN00760947 | Crude Extract | 31.3 |
| 5 | 25641 | Porifera | SN00731867 | Crude Extract | 250.0 |
| 6^†^ | 26051 | Porifera | SN00731005 | Crude Extract | 62.5 |
| 7 | 25642 | Porifera | SN00731868 | Crude Extract | 125.0 |
| 8 | 24132 | Porifera | SN00734102 | Crude Extract | 125.0 |
| 9 | 19033 | Porifera | SN00733107 | MeOH Eluent | 62.5 |
| 10^†^ | 19033 | Porifera | SN00733110 | Crude Extract | 31.3 |
| 11 | 19039 | Porifera | SN00733134 | Crude Extract | 125.0 |
| 12 | 19039 | Porifera | SN00733131 | MeOH Eluent | 250.0 |
| 13^†^ | 20608 | Porifera | SN00760956 | 75% MeOH Eluent | 31.3 |
| 14^†^ | 20608 | Porifera | SN00760958 | MeOH Eluent | 62.5 |
| 15 | 25691 | Porifera | SN00732374 | Crude Extract | 125.0 |
| 16^†^ | 24307 | Porifera | SN00730755 | 75% MeOH Eluent | 31.3 |
| 17 | 24307 | Porifera | SN00730707 | Crude Extract | 500.0 |
| 18 | 25663 | Chordata | SN00732222 | Crude Extract | 15.6 |
| 19 | 25663 | Chordata | SN00732228 | 75% MeOH Eluent | 125.0 |
| 20 | 26104 | Porifera | SN00734298 | Crude Extract | 62.5 |
| 21 | 25658 | Porifera | SN00732162 | Crude Extract | 125.0 |
| 22 | 25658 | Porifera | SN00732159 | MeOH Eluent | 500.0 |
| 23 | 24348 | Echinodermata | SN00739901 | 30% MeOH Eluent | 500.0 |

^†^ Samples with promising biological activity that were subjected to further investigation.

**Table S2. Biological Origin of Samples of Interest.**

| Entry^†^ | AIMS Sample Code | Biological origin | QCL Sample Number | Fraction |
| --- | --- | --- | --- | --- |
| 4 | 20608 | *Lendenfeldia sp.* | SN00760947 | crude extract |
| 6 | 26051 | *Ircinia gigantea* | SN00731005 | crude extract |
| 10 | 19033 | *Dysidea herbacea* | SN00733110 | crude extract |
| 13 | 20608 | *Lendenfeldia sp.* | SN00760956 | 75% MeOH eluent |
| 14 | 20608 | *Lendenfeldia sp.* | SN00760958 | 100% MeOH eluent |
| 16 | 24307 | Class Demospongiae^‡^ | SN00730755 | 75% MeOH eluent |

^†^ Corresponds to Entry Number in Table S1 above.

^‡^ More detailed taxonomic classification not available.

**Table S3. Cytotoxicity of the 23 most active extracts against HepG2, HEK 293, THP-1 and A549 cell lines.**

| Entry | AIMS Sample Code | QCL Sample Number | Cytotoxicity (% cell survival)^†^ | | | |
| --- | --- | --- | --- | --- | --- | --- |
|  |  |  | **THP-1** | **A549** | **HEK** | **HepG2** |
| 1 | 22565 | SN00739718 | 19 | 99 | 97 | >100 |
| 2 | 24606 | SN00732793 | 53 | 47 | 96 | 98 |
| 3 | 24307 | SN00731867 | 202 | 113 | 97 | 95 |
| 4 | 20608 | SN00731005 | 214 | 110 | 98 | 97 |
| 5 | 25641 | SN00731868 | 66 | 106 | 109 | 99 |
| 6 | 26051 | SN00734102 | 23 | 102 | 110 | >100 |
| 7 | 25642 | SN00732374 | 74 | 55 | 92 | 99 |
| 8 | 24132 | SN00732162 | 21 | 46 | 48 | 96 |
| 9 | 19033 | SN00730707 | 194 | 111 | 103 | 93 |
| 10 | 19033 | SN00733110 | 87 | 91 | 98 | 91 |
| 11 | 19039 | SN00733107 | 15 | 6 | 4 | 96 |
| 12 | 19039 | SN00733134 | 18 | 5 | 4 | >100 |
| 13 | 20608 | SN00733131 | 42 | 96 | 102 | 100 |
| 14 | 20608 | SN00760956 | 173 | 106 | 95 | 98 |
| 15 | 25691 | SN00760958 | 162 | 108 | 95 | 96 |
| 16 | 24307 | SN00760947 | 169 | 101 | 102 | 100 |
| 17 | 24307 | SN00730755 | 168 | 106 | 96 | >100 |
| 18 | 25663 | SN00730771 | 30 | 101 | 104 | 100 |
| 19 | 25663 | SN00732222 | 19 | 89 | 97 | 100 |
| 20 | 26104 | SN00732228 | 19 | 88 | 105 | 98 |
| 21 | 25658 | SN00732162 | 53 | 105 | 105 | 102 |
| 22 | 25658 | SN00732159 | 194 | 108 | 98 | 87 |
| 23 | 24348 | SN00739901 | 31 | 110 | 94 | >100 |

^†^ Cells (2 × 10^5^ cells/well) were seeded and incubated with 50 µg mL^−1^ of extract samples for 4 days under humidified incubation at 37 °C with 5% CO_2_.

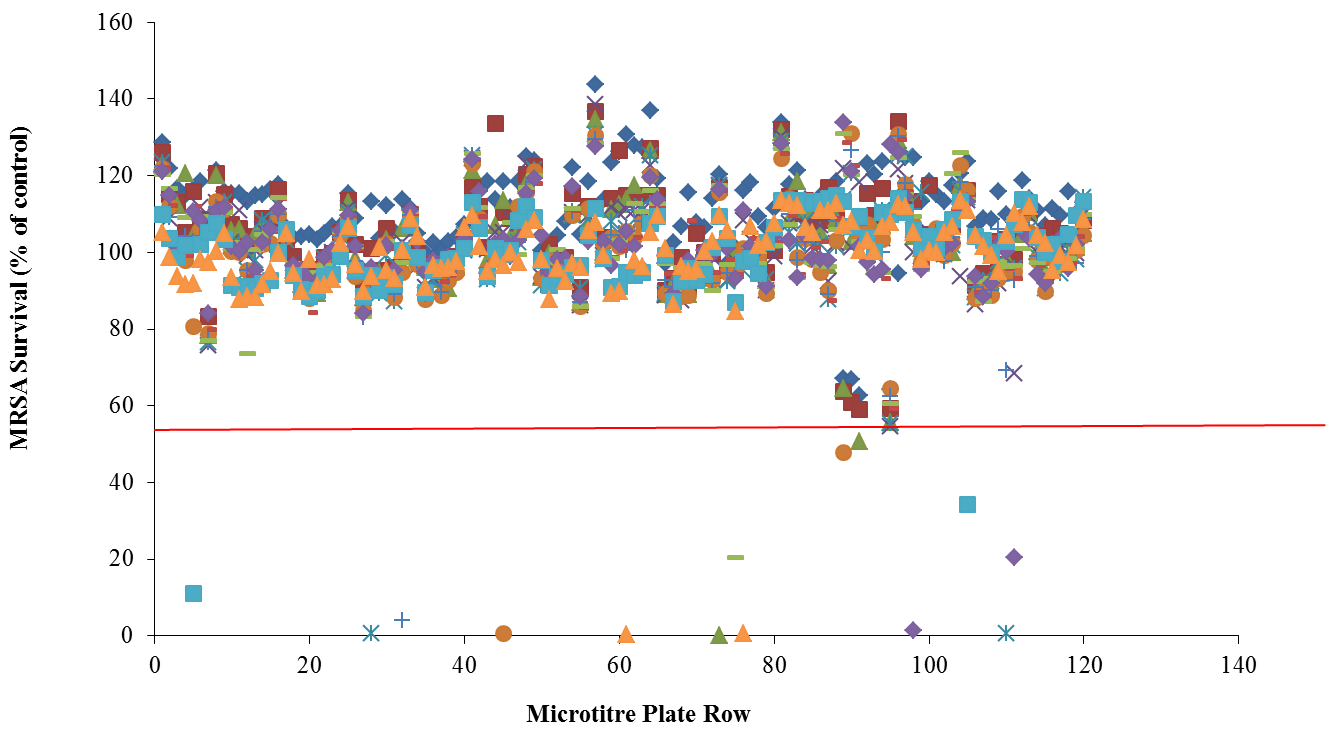

**Figure S1. Primary screen of marine natural product extracts for activity against MRSA.** MRSA (OD_600_ =0.001) was incubated with each of the 1434 library samples at a final concentration of 50 µg mL^−1^, under optimum assay conditions as described in the Experimental section of the manuscript. A cut-off of less than 50% MRSA survival was chosen to represent positive hits (shown as solid red line). The figure displays the mean survival ± SD for triplicate samples and represents data from three independent repeats.

### 2. HPLC FRACTIONATION AND PURIFICATION

**2.1.	Summary of HPLC Methods.**

**Table S4**. **Summary of HPLC methods for the purification of key samples of interest.**

| AIMS  Sample Code | Source Information | HPLC Step 1 | HPLC Step 2 |
| --- | --- | --- | --- |
| 19033 | Phylum: *Porifera*  Class: *Demospongiae*  Order: *Dendroceratida* Family: *Dysideidae*  Genus: *Dysidea*  Species: *herbacea* | Reversed-Phase (C18), 1% TFA in  H_2_O/MeCN, 80 min 0-100% MeCN,  12 mL min^-1^ | Reversed-Phase (C18), 1% TFA in H_2_O/MeCN,  20 min 0-40% MeCN, 60 min 40-100% MeCN, 10 min 100% MeCN, 7 mL min^-1^ |
| 20608 | Phylum: *Porifera*  Class: *Demospongiae*  Order: *Dictyoceratida*  Family: *Spongiidae*  Genus: *Lendenfeldia* | Reversed-Phase (C18), 1% TFA in  H_2_O/MeCN, 80 min 0-100% MeCN,  12 mL min^-1^ | Reversed-Phase (C18), 1% TFA in H_2_O/MeCN,  20 min 0-40% MeCN, 60 min 40-100% MeCN, 10 min 100% MeCN, 7 mL min^-1^ |
| 26051 | Phylum: *Porifera*  Class: *Demospongiae*  Order: *Dictyoceratida*  Family: *Irciniidae*  Genus: *Ircinia* | Reversed-Phase (C18), 1% TFA in  H_2_O/MeCN, 80 min 0-100% MeCN,  12 mL min^-1^ | Reversed-Phase (C18), 1% TFA in H_2_O/MeCN,  20 min 0-40% MeCN, 60 min 40-100% MeCN, 10 min 100% MeCN, 7 mL min^-1^ |

**2.2.	Fractionation of AIMS Sample 19033.**

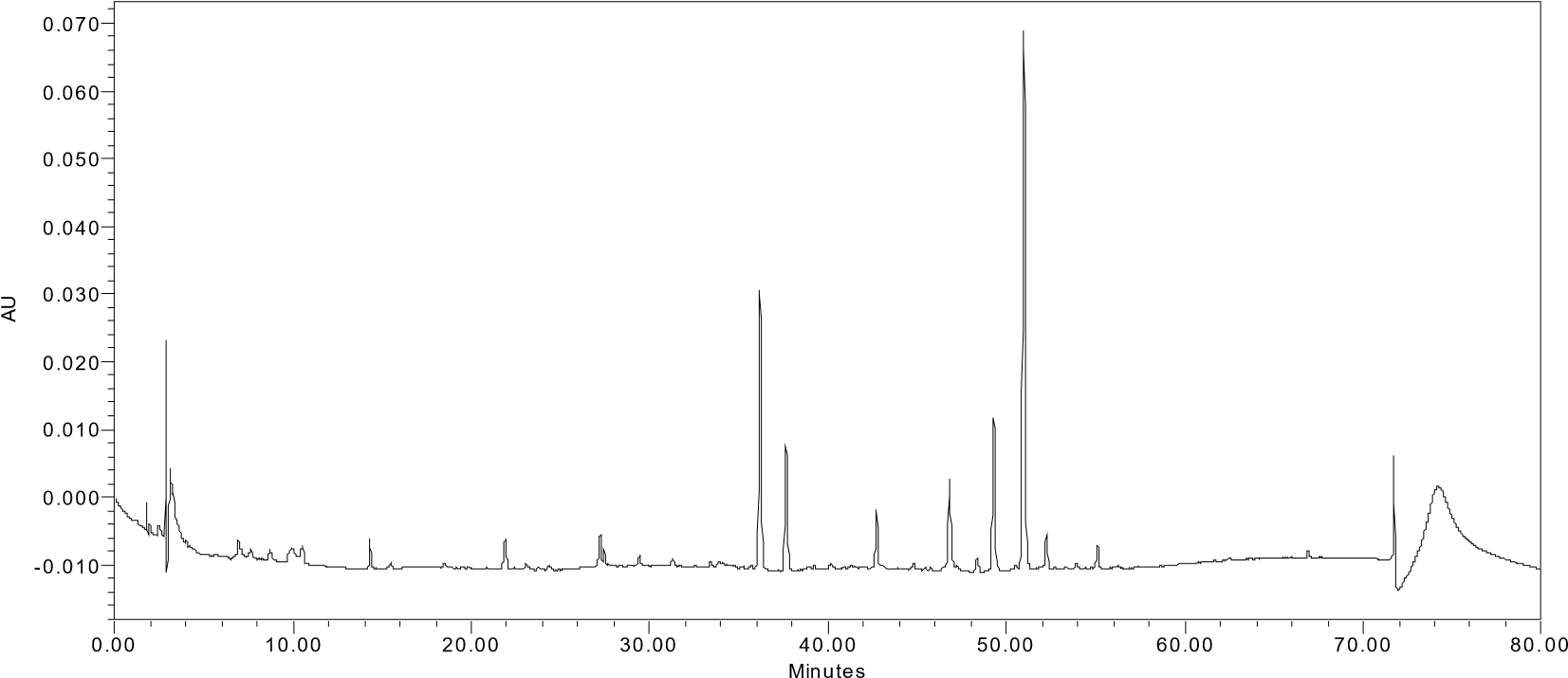

**Figure S2.** **Representative HPLC trace for sample 19033.** Preparative HPLC was carried out using a C18 RP-HPLC column and a gradient with 0 to 100 % acetonitrile-H_2_O, flow rate 1 mL min^-1^, monitored at 254 nm, to yield 80 fractions over 80 minutes.

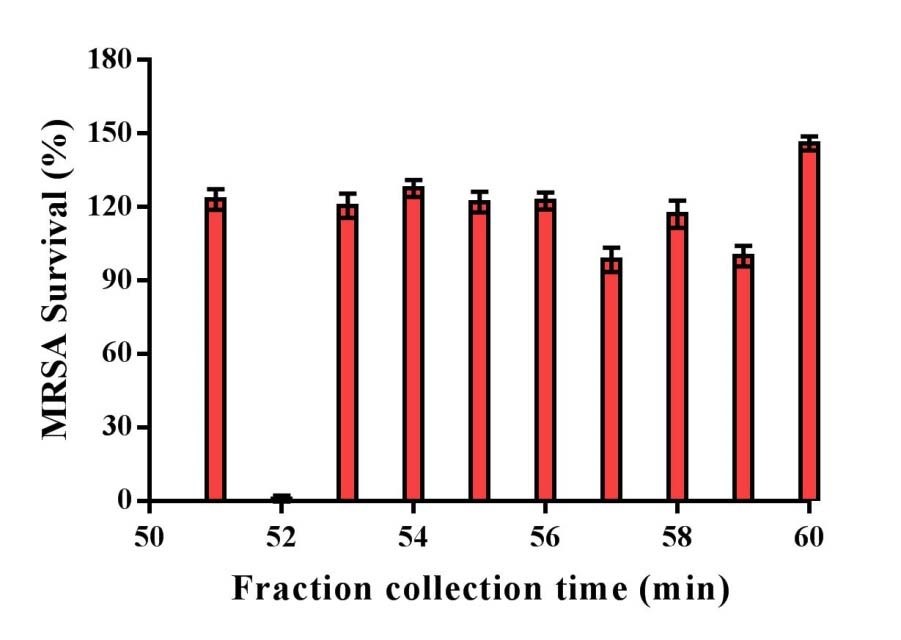

**Figure S3.** **Bioassay of selected fractions from HPLC purification of sample 19033.** Fractions were tested for bioactivity against MRSA. Data show mean survival ± SEM of duplicate samples.

**2.3.	Fractionation of AIMS Sample 20608.**

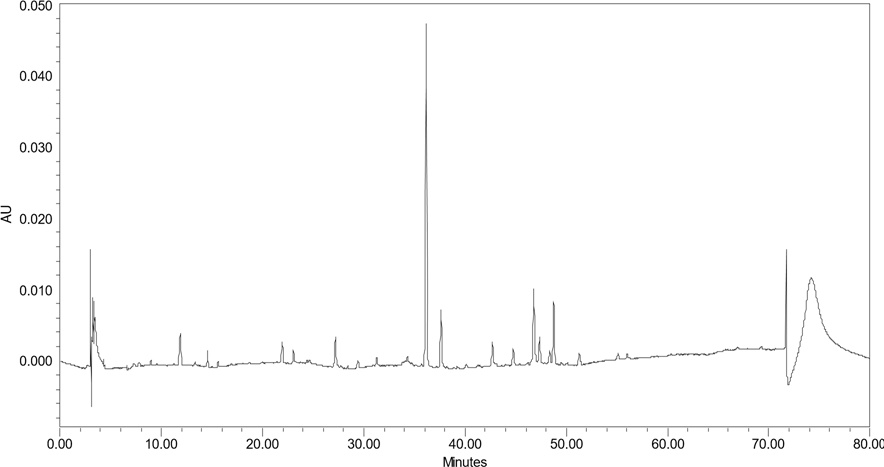

**Figure S4. Representative HPLC trace for sample 20608.** Preparative HPLC was carried out using a C18 RP-HPLC column and a gradient with 0 to 100 acetonitrile-H_2_O, flow rate 1 mL
min^-1^, monitored at 254 nm, to yield 80 fractions over 80 minutes.

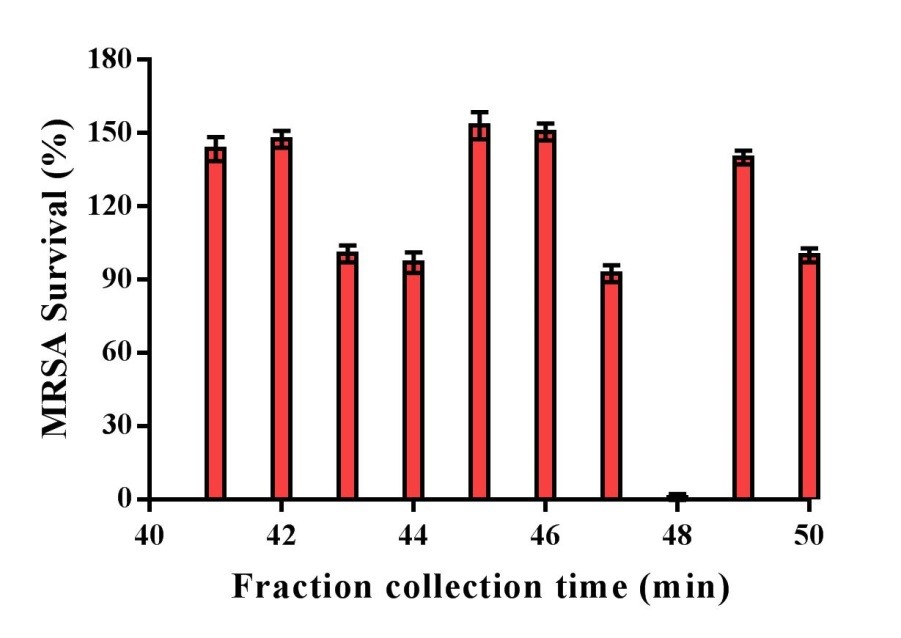

**Figure S5. Bioassay of selected fractions from HPLC purification of sample 20608.** Fractions were tested for bioactivity against MRSA. Data show mean survival ± SEM of duplicate samples.

2.4. **Fractionation of AIMS Sample 26051.**

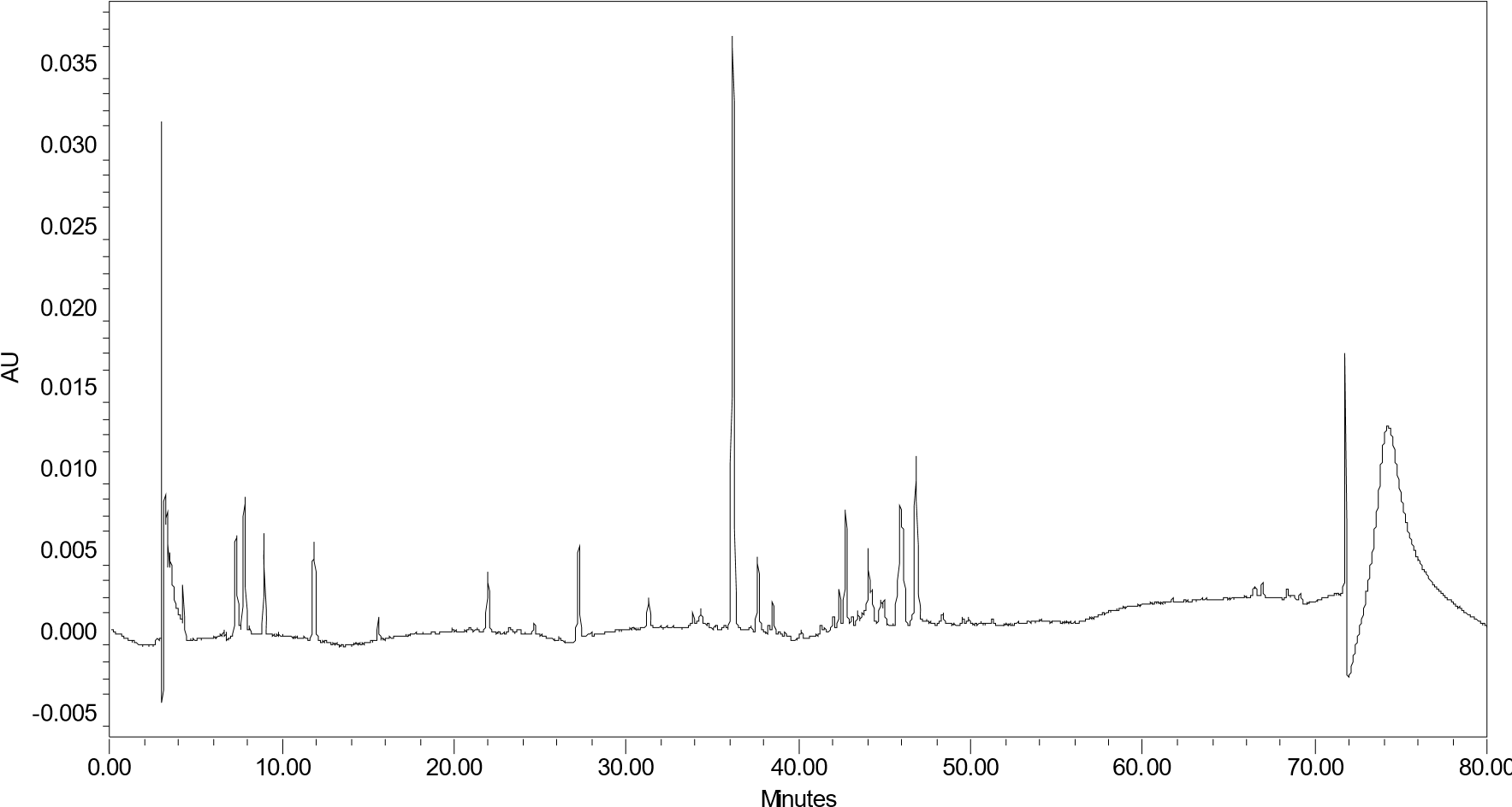

**Figure S6. Representative HPLC trace for sample 26051.** Preparative HPLC was carried out using a C18 RP-HPLC column and a gradient with 0 to 100 acetonitrile-H_2_O, flow rate 1 mL
min^-1^, monitored at 254 nm, to yield 80 fractions over 80 minutes.

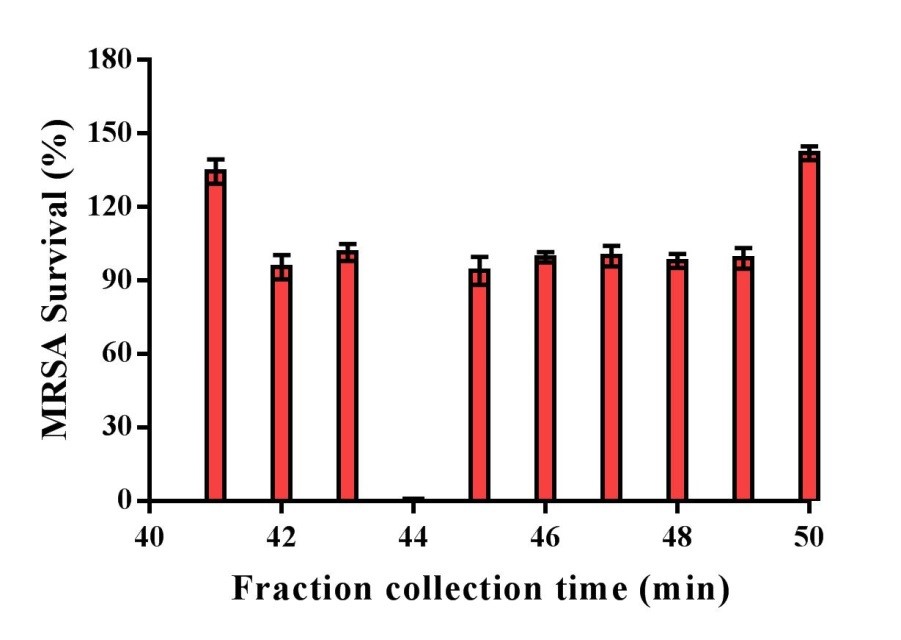

**Figure S7. Bioassay of selected fractions from HPLC purification of sample 26051.** Fractions were tested for bioactivity against MRSA. Data show mean survival ± SEM of duplicate samples.

### 3. MASS SPECTROMETRY DATA FOR NATURAL PRODUCTS.

**3.1	MS Analysis of AIMS Samples 19033.**

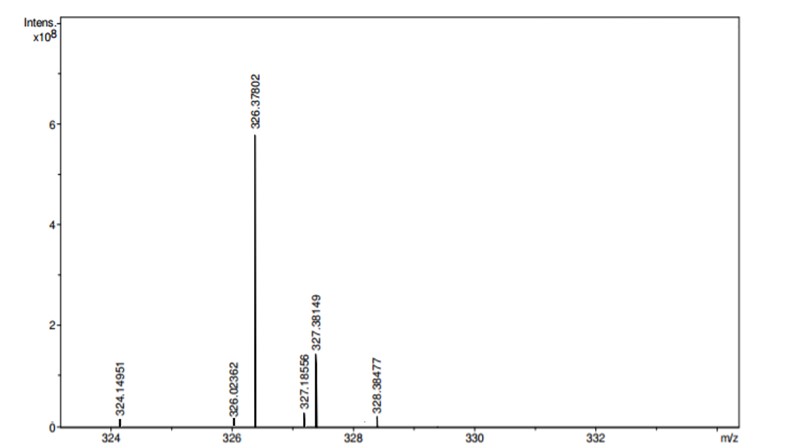

**Figure S8. HRMS spectrum of the biologically active component of extract 19033.**

MS analysis of the bioactive component of sample number 19033 shows a molecular ion at experimental *m/z* 326.37813; assuming [CHNO+Na_0-1_]^+^ theoretical *m/z* for [C_22_H_48_N]^+^ = 326.37802; ∆m = 0.11 ppm, RDBE = 0 (Figure S8). The MS/MS fragmentation shows fragments consistent with saturated hydrocarbon chains, up to C_10_ in length, leading to the proposal of the tertiary amine structure **3** for this compound. Key daughter ions and suggested fragment structures are represented in Table S5 and fragmentation pathways in Figure S9.

**Table S5. MS/MS**, MS^3^ and MS^4^ **data and assignments for AIMS samples 19033.**

| Parent ion peak | MS/MS or MS^3^ pattern | Neutral loss (*m/z* or amu) | Daughter ion peak | Fragment form | Predictive structure |
| --- | --- | --- | --- | --- | --- |
| 326 [C_22_H_48_ N+H]^+^  (rdbe 0) | → 324 | [M-2H] | 324 | [C_22_H_46_N]^+^ | Double bond or ring formation |
|  | → 241 | (C_15_H_39_N) or  (C_14_H_37_) | 107 | [C_7_H_9_N]^+^ or  [C_8_H_11]_^+^ | 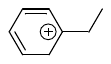or 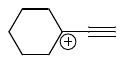 |
|  |  | (C_12_H_23_) | 159 | [C_10_H_25_]^+^ | C_10_ hydrocarbon chain |
|  | → 241→159 | (C_17_H_43_N) or  (C_18_H_45_) | 65 | [C_5_H_5_]^+^ or  [C_4_H_3_N]^+^ | 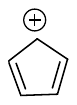 or 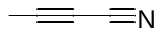 |
|  | → 243 | (C_6_H_11_) | 243 | [C_16_H_37_N]^+^ | 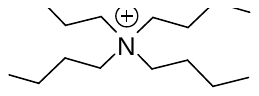 |
|  | → 186 | (C_18_H_39_N) or  (C_19_H_37_) | 57 | [C_4_H_9_]^+^ or  [C_3_H_7_N]^+^ | 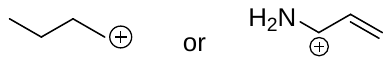 |
|  |  | (C_17_H_37_N) or  (C_18_H_39_) | 71 | [C_5_H_11_]^+^ | 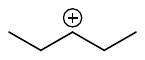 |
|  |  | (C_16_H_35_N) or  (C_17_H_37_) | 85 | [C_6_H_13_]^+^ or  [C_5_H_11_N]^+^ | 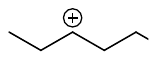 |
|  |  | (C_15_H_33_N) or  (C_16_H_35_) | 99 | [C_7_H_15_]^+^ or  [C_6_H_13_N]^+^ | 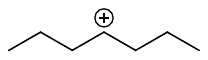 |
|  | → 186 → 85 | (C_18_H_39_N) or  (C_19_H_41_) | 57 | [C_4_H_9_]^+^ or  [C_3_H_7_N]^+^ | 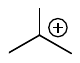 |
|  |  | (C_18_H_41_N) or  (C_19_H_43_) | 55 | [C_4_H_7_]^+^ or  [C_3_H_5_N]^+^ | 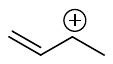 |

rdbe = ring or double bond equivalents

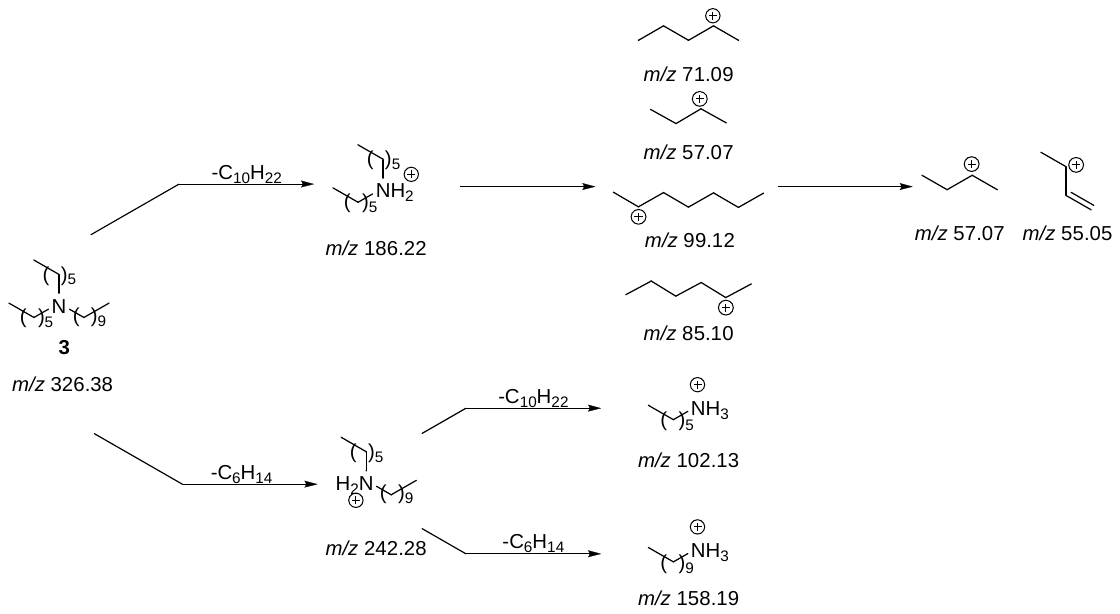

**Figure S9. Key steps in the proposed MS/MS**, MS^3^ and MS^4^ **fragmentation pathway of the active component of AIMS sample 19033.**

**3.2	MS Analysis of AIMS Sample 20608**.

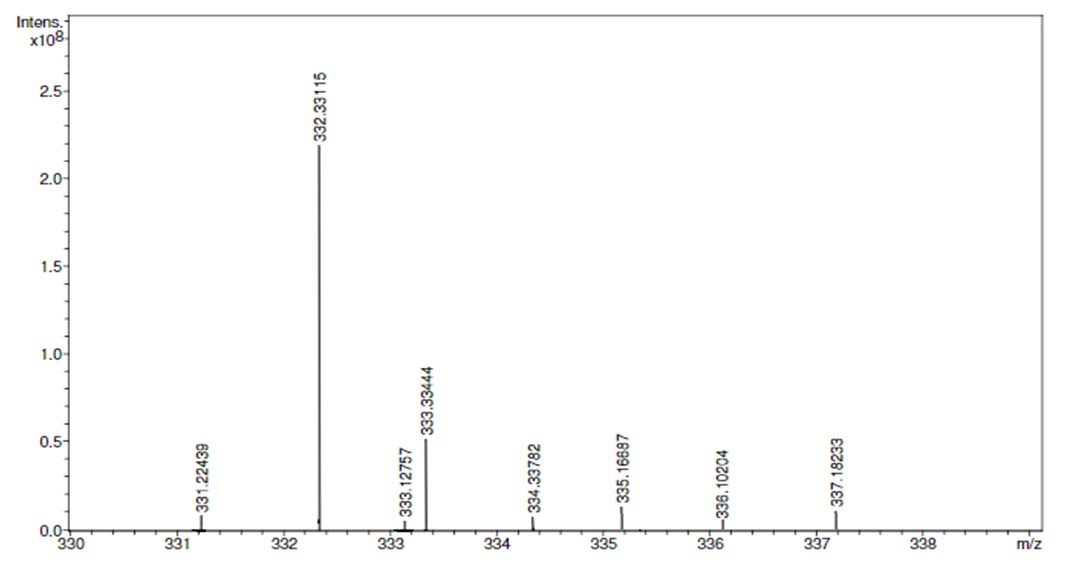

**Figure S10. High resolution mass spectrum of the biologically active component of AIMS extract 20608 (QCL samples SN00760947, SN00760956 and SN00760958).**

By MS (Figure S10), accurate mass measurements for [M+H]^+^ at *m/z* 332 (the strongest signal in this spectrum) returned an experimental accurate mass of 332.33115; theoretical (assuming [CHNO+Na_0-1_]^+^) for C_23_H_42_N = 332.33118 (Δm = 0.03 ppm). No evidence of deuterium incorporation was observed (Table S6, isotopic fine structure around 333.33 *m/z* (data not shown)). MS/MS data revealed key fragment ions corresponding to neutral loss of C_7_H_8_ (rdbe = 4, potentially toluene) to form [C_16_H_34_N]^+^ (rdbe = 0). Neutral loss of [C_7_H_7_] suggests the presence of tropylium ion, a further indication of the presence of a tolyl moiety. MS/MS measurements of the fragment at *m/z* 259 showed fragmentation to [C_9_H_7_]^+^ potentially 1-ethynyl-4-methylbenzene or 1H-inden-2-ylium (Figure S11). Taken together, these data suggest that the compound is a tertiary amine in which one substituent is either a methylphenyl group, with the methyl group in an undetermined position (*ortho*, *meta* or *para*), or a benzyl group. The other two substituent are saturated hydrocarbon chains, including a total of 16 carbon atoms. One of these chains may be as long as C_10_ (based on the largest fragments), with some branching on one or both chains likely present. Thus the generalised structures **1** and **2** were proposed for the active agents in AIMS sample 20608.

| Parent ion peak | MS/MS or MS^3^ pattern | Neutral loss (*m/z* or amu) | Daughter ion  peak | Fragment form | | Predictive structure |
| --- | --- | --- | --- | --- | --- | --- |
| 332  [C_23_H_42_N+H]^+^ (rdbe 4) |  | (C_7_H_10_) | 238 | [C_16_H_32_N]^+^ | (rdbe 0) | 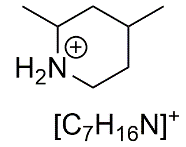 |
|  | → 240  [C_16_H_34_N]^+^  loss of (C_7_H_8_) | (C_11_H_25_N) | 69 | [C_5_H_9_]^+^ | (rdbe 1) |  |
|  |  | (C_10_H_23_N) | 83 | [C_6_H_11_]^+^ | (rdbe 1) |  |
|  |  | (C_9_H_21_N) | 97 | [C_7_H_13_]^+^ | (rdbe 1) |  |
|  |  | (C_8_H_19_N) | 111 | [C_8_H_15_]^+^ | (rdbe 1) |  |
|  |  | (C_9_H_18_) | 114 | [C_7_H_16_N]^+^ | (rdbe 1) |  |
|  | → 240 → 97 | (C_3_H_6_) | 55 | [C_4_H_7_]^+^ | (rdbe 1) |  |
|  | → 240 → 114 | (C_10_H_20_) | 100 | [C_6_H_14_N]^+^ | (rdbe 1) |  |
|  | → 259 | (C_10_H_24_) | 115 | [C_9_H_7_]^+^ | (rdbe 6) | 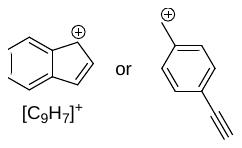 |
|  |  | (C_4_H_10_) | 201 | [C_15_H_21_]^+^ | (rdbe 5) |  |
|  |  | (C_2_H_12_) or  [M – C_3_] | 223 | [C_17_H_19_]^+^  or  [C_16_H_31_]^+^ | (rdbe 8)  or  (rdbe 1) |  |
|  |  | [M – 2H] | 257 | [C_19_H_29_]^+^ | (rdbe 6) |  |
|  |  | (C_16_H_34_N) | 91 | [C_7_H_7_]^+^ | (rdbe 5) | 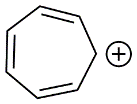 |

**Table S6. MS/MS**, MS^3^ and MS^4^ **data and assignments for AIMS sample 20608.**

rdbe = ring or double bond equivalents

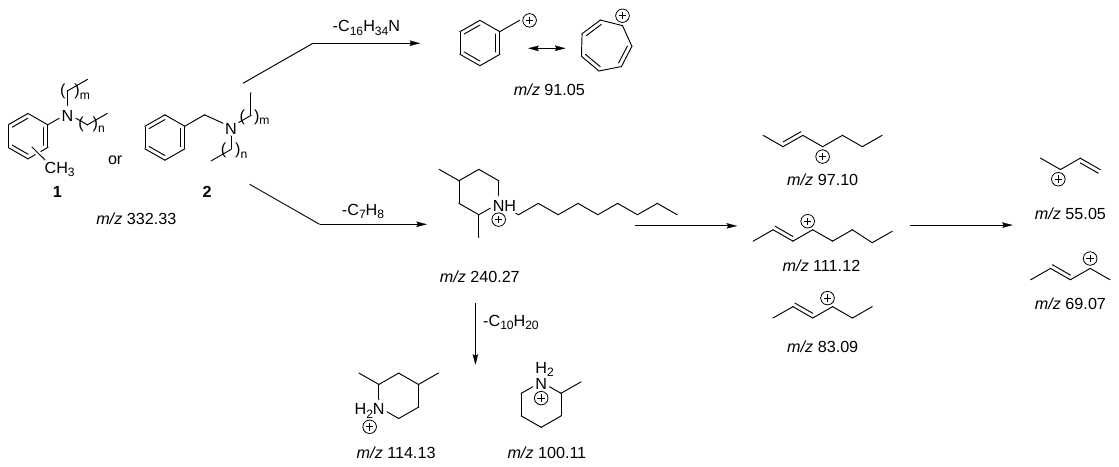

**Figure S11. Proposed MS/MS**, MS^3^ and MS^4^ **fragmentations for compound 1/2 from AIMS sample 20608.**

**3.3	MS Analysis of AIMS Sample 26051.**

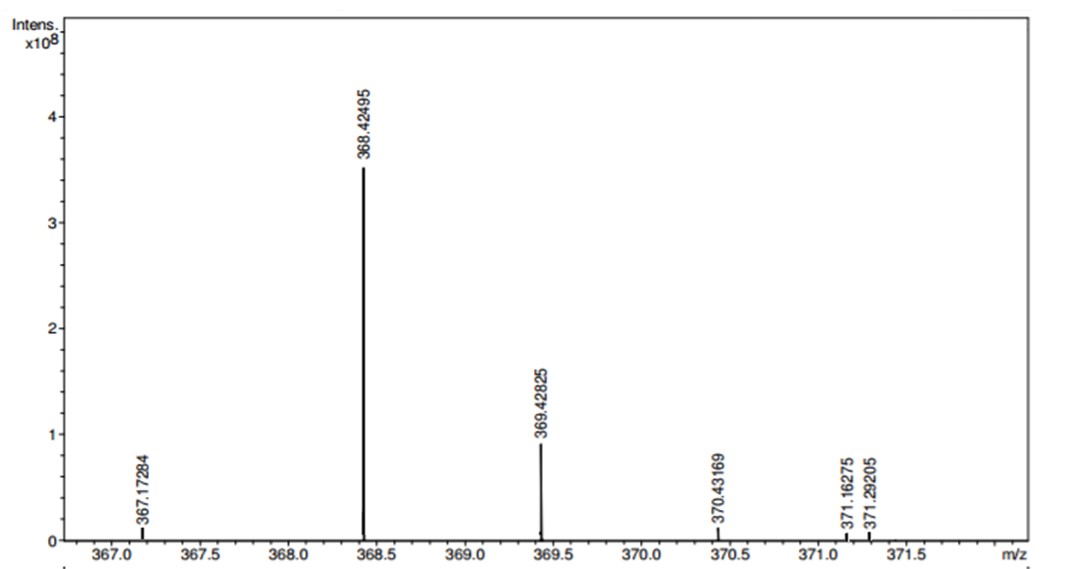

**Figure S12. High resolution mass spectrum for the biologically active component of AIMS extract 26051.**

Fractionation of the sample 26051 (Table S4 and Figure S4) afforded an active component of accurate MS (Figure S12) of experimental mass 368.42508; theoretical mass (assuming [CHNO+Na_0-1_]^+^) for C_25_H_54_N = 368.42495; ∆m = 0.13 ppm, RDBE = 0. MS/MS spectra showed five daughter ions giving fragments consistent with the presence of long hydrocarbon chains (Table S7), suggesting that this compound is fully saturated (0 RDBE) and is either a tertiary amine or quaternary amine salt, with one propyl chain and one of the other chains at least C_10_ in length (Figure S13).

**Table S7. MS/MS** and MS^3^ **data and assignments for AIMS samples 26051.**

| **Parent ion** | **MS/MS** or MS^3^ **pattern** | **Neutral loss (*m/z* or amu)** | **Daughter ion peak** | **Fragment form** | | **Predictive structure** |
| --- | --- | --- | --- | --- | --- | --- |
| **368**  **[C_25_H_54_N+H]^+^ (rdbe 0)** |  | C_3_H_8_ | 324 | [C_22_H_46_N]^+^ | (1 rdbe) | Long hydrocarbon chain |
|  |  | M-3H | 365 | [C_26_H_53_]^+^  or  [C_25_H_49_O]^+^ |  |  |
|  |  | NH_2_+H_2_ | 349 | [C_25_H_49_]^+^ | (1 rdbe) |  |
|  |  | C_3_H_9_N | 309 | [C_22_H_45_]^+^ | (0 rdbe) |  |
|  |  | C_5_H_15_N | 279 | [C_20_H_39_]^+^ | (1 rdbe) |  |
|  | → 324 | C_3_H_7_N | 267 | [C_19_H_39_]^+^ | (0 rdbe) |  |
|  |  | C_12_H_29_N | 137 | [C_10_H_17_]^+^ | (2 rdbe) |  |
|  |  | C_13_H_27_N | 123 | [C_9_H_15_]^+^ | (2 rdbe) |  |
|  |  | C_14_H_25_N | 109 | [C_8_H_13_]^+^ | (2 rdbe) |  |
|  |  | C_15_H_25_N | 97 | [C_7_H_13_]^+^ | (1 rdbe) |  |
|  | → 309 | C_4_H_10_ | 251 | [C_18_H_35_]^+^ | (1 rdbe) |  |

rdbe = ring or double bond equivalents

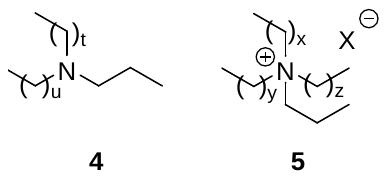

**Figure S13. Proposed general structures for the tertiary amines isolated from 26051; (t + u) = 20; (x + y + z) = 1.**

### 4. MASS SPECTROMETRY DATA FOR SYNTHETIC COMPOUNDS.

**Table S8. MS/MS**, MS^3^, MS^4^ and MS^5^ **data and assignments for synthetic compound 1.**

| **Parent ion peak** | **MS/MS, MS^3^ or MS^4^ pattern** | **Neutral loss (m/z or amu)** | **Daughter ion Peak** | | **Fragment form** | |
| --- | --- | --- | --- | --- | --- | --- |
| **332.3**  **(C_23_H_42_N^+^)** |  | (C_7_H_8_) | | 240.3 | [C_16_H_34_N]^+^ | (rdbe 1) |
|  |  | (C_8_H_16_) | | 220.2 | [C_15_H_26_N]^+^ | (rdbe 4) |
|  |  | (C_16_H_32_) | | 108.1 | [C_7_H_10_N]^+^ | (rdbe 4) |
|  |  | (C_7_H_16_) | | 232.2 | [C_16_H_26_N]^+^ | (rdbe 5) |
|  |  | (C_6_H_13_N) | | 233.2 | [C_17_H_29_]^+^ | (rdbe 4) |
|  | → 240 (C_16_H_34_N^+^) | (C_14_H_22_) | | 142.2 | [C_9_H_20_N]^+^ | (rdbe 1) |
|  |  | (C_15_H_24_) | | 128.2 | [C_8_H_18_N]^+^ | (rdbe 1) |
|  |  | (C_18_H_33_N) | | 69.1 | [C_5_H_9_]^+^ | (rdbe 2) |
|  | → 233 (C_17_H_29_^+^) | (C_7_H_16_) | | 232.2 | [C_16_H_26_N]^+^ | (rdbe 5) |
|  |  | (C_13_H_27_N) | | 135.1 | [C_10_H_15_]+ | (rdbe 4) |
|  |  | (C_14_H_30_) | | 134.1 | [C_9_H_12_N]^+^ | (rdbe 4) |
|  | → 220 (C_15_H_26_N^+^) | (C_15_H_27_N) | | 93.1 | [C_7_H_9_]^+^ | (rdbe 4) |
|  |  | (C_18_H_33_N) | | 69.1 | [C_5_H_9_]^+^ | (rdbe 2) |
|  |  | (C_16_H_32_) | | 108.1 | [C_7_H_10_N]^+^ | (rdbe 4) |
|  |  | (C_15_H_32_) | | 120.1 | [C_8_H_10_N]^+^ | (rdbe 5) |
|  |  | (C_18_H_31_N) | | 71.1 | [C_5_H_11_]^+^ | (rdbe 1) |
|  |  | (C_9_H_18_) | | 206.2 | [C_14_H_24_N]^+^ | (rdbe 4) |
|  |  | (C_16_H_35_N) | | 91.1 | [C_7_H_7_]^+^ | (rdbe 5) |
|  | → 240 → 142.2 | (C_18_H_33_N) | | 69.1 | [C_5_H_9_]^+^ | (rdbe 2) |
|  |  | (C_21_H_36_) | | 44.1 | [C_2_H_6_N]^+^ | (rdbe 1) |
|  | → 240 → 142 → 128 | (C_15_H_27_N) | | 111.1 | [C_8_H_15_]^+^ | (rdbe 2) |
|  |  | (C_18_H_33_N) | | 69.1 | [C_5_H_9_]^+^ | (rdbe 2) |
|  |  | (C_19_H_33_N) | | 57.1 | [C_4_H_9_]^+^ | (rdbe 1) |
|  | → 240 → 220 → 108 | (C_16_H_33_N) | 93.1 | | [C_7_H_9_]^+^ | (rdbe 4) |

rdbe = ring or double bond equivalents

**Table S9. MS/MS data and assignments for synthetic compound 2.**

| **Parent ion peak** | **MS/MS pattern** | **Neutral loss (m/z or amu)** | **Daughter ion Peak** | **Fragment form** | |
| --- | --- | --- | --- | --- | --- |
| **332.3**  **(C_23_H_42_N^+^)** |  | (C_7_H_8_) | 240.3 | [C_16_H_34_N]^+^ | (rdbe 1) |
|  |  | (C_14_H_22_) | 142.2 | [C_9_H_20_N]^+^ | (rdbe 1) |
|  |  | (C_16_H_35_N) | 91.1 | [C_7_H_7_]^+^ | (rdbe 5) |
|  | → 240.3 (C_16_H_34_N^+^) | (C_14_H_2_2) | 142.2 | [C_9_H_20_N]^+^ | (rdbe 1) |
|  |  | (C_15_H_24_) | 128.2 | [C_8_H_18_N]^+^ | (rdbe 1) |
|  |  | (C_18_H_33_N) | 69.1 | [C_5_H_9_]^+^ | (rdbe 2) |
|  | → 240 → 142.2 | (C_15_H_27_N) | 111.1 | [C_8_H_15_]^+^ | (rdbe 2) |
|  |  | (C_18_H_33_N) | 69.1 | [C_5_H_9_]^+^ | (rdbe 2) |
|  | → 240 → 142 → 128 | (C_15_H_27_N) | 111.1 | [C_8_H_15_]^+^ | (rdbe 2) |
|  |  | (C_18_H_33_N) | 69.1 | [C_5_H_9_]^+^ | (rdbe 2) |

rdbe = ring or double bond equivalents

**Table S10. MS/MS data and assignments for synthetic compound 3.**

| **Parent ion peak** | **MS/MS pattern** | **Neutral loss (m/z or amu)** | **Daughter ion Peak** | **Fragment form** | |
| --- | --- | --- | --- | --- | --- |
| **326.4 (C_22_H_48_N^+^)** |  | (C_6_H_12_) | 242.3 | [C_16_H_36_N]^+^ | (rdbe 0) |
|  |  | (C_10_H_20_) | 186.2 | [C_12_H_28_N]^+^ | (rdbe 0) |
|  |  | (C_12_H_24_) | 158.2 | [C_10_H_24_N]^+^ | (rdbe 0) |
|  |  | (C_16_H_32_) | 102.1 | [C_6_H_16_N]^+^ | (rdbe 0) |
|  | →242.3 (C_16_H_36_N^+^) | (C_6_H_14_) | 240.3 | [C_16_H_34_N]^+^ | (rdbe 1) |
|  |  | (C_12_H_24_) | 158.2 | [C_10_H_24_N]^+^ | (rdbe 0) |
|  |  | (C_16_H_32_) | 102.1 | [C_6_H_16_N]^+^ | (rdbe 0) |
|  |  | (C_18_H_39_N) | 85.1 | [C_6_H_13_]^+^ | (rdbe 1) |
|  |  | (C_17_H_37_N) | 71.1 | [C_5_H_11_]^+^ | (rdbe 1) |
|  | →186.2 (C_12_H_28_N^+^) | (C_16_H_32_) | 102.1 | [C_6_H_16_N]^+^ | (rdbe 0) |
|  |  | (C_16_H_35_N) | 85.1 | [C_6_H_13_]^+^ | (rdbe 1) |
|  |  | (C_18_H_39_N) | 57.1 | [C_4_H_9_]^+^ | (rdbe 1) |
|  | →102.1 (C_6_H_16_N^+^) | (C_16_H_35_N) | 85.1 | [C_6_H_13_]^+^ | (rdbe 1) |
|  |  | (C_18_H_39_N) | 57.1 | [C_4_H_9_]^+^ | (rdbe 1) |
|  | → 158.2 (C_10_H_24_N^+^) | (C_16_H_35_N) | 85.1 | [C_6_H_13_]^+^ | (rdbe 1) |
|  |  | (C_18_H_39_N) | 57.1 | [C_4_H_9_]^+^ | (rdbe 1) |
|  |  | (C_17_H_37_N) | 71.1 | [C_5_H_11_]^+^ | (rdbe 1) |
|  |  | (C_12_H_26_) | 156.2 | [C_10_H_22_N]^+^ | (rdbe 1) |
|  | → 158.2 → 85.1 | (C_19_H_41_N) | 43.1 | [C_3_H_7_]^+^ | (rdbe 0) |
|  |  | (C_18_H_39_N) | 57.1 | [C_4_H_9_]^+^ | (rdbe 1) |

rdbe = ring or double bond equivalents

**Table S11.** **MS/MS data and assignments for synthetic compound 13**.

| **Parent ion peak** | **MS/MS pattern** | **Neutral loss (m/z or amu)** | **Daughter ion Peak** | **Fragment form** | |
| --- | --- | --- | --- | --- | --- |
| **354.4**  **(C_24_H_52_ N^+^)** |  | (C_8_H_16_) | 242.3 | [C_16_H_36_N]^+^ | (rdbe 0) |
|  |  | (C_16_H_32_) | 130.2 | [C_8_H_20_N]^+^ | (rdbe 0) |
|  | → 242.3 (C_16_H_36_N^+^) | (C_19_H_41_N) | 71.1 | [C_5_H_11_]^+^ | (rdbe 1) |
|  |  | (C_16_H_32_) | 130.2 | [C_8_H_20_N]^+^ | (rdbe 0) |
|  | → 242.3 → 71.1 | (C_21_H_45_N) | 43.1 | [C_3_H_7_]^+^ | (rdbe 0) |
|  | → 242.3 → 130.2 | (C_16_H_34_) | 128.2 | [C_8_H_18_N]^+^ | (rdbe1) |
|  |  | (C_19_H_41_N) | 71.1 | [C_5_H_11_]^+^ | (rdbe 1) |
|  |  | (C_21_H_45_N) | 43.1 | [C_3_H_7_]^+^ | (rdbe 0) |
|  |  | (C_20_H_43_N) | 57.1 | [C_4_H_9_]^+^ | (rdbe 1) |

rdbe = ring or double bond equivalents
